# Predicting porcini: a decade of sporocarp monitoring reveals the meteorological triggers of *Boletus edulis* fruiting in central European beech forests

**DOI:** 10.64898/2025.12.12.693895

**Authors:** Etienne Brejon Lamartiniere, Joseph Ivan Hoffman

## Abstract

Despite recent molecular and genomic advances, the reproductive biology of ectomycorrhizal fungi (EMF) remains poorly understood. In particular, the meteorological cues triggering sporocarp formation remain largely unknown, limiting our ability to induce fruiting in the laboratory. To address this knowledge gap, we analysed a decade-long (2015–2024) dataset of daily, near-exhaustive sporocarp observations from an intensively monitored population of *Boletus edulis* in a central European beech forest near Bielefeld, Germany. Using generalised linear mixed-effect models, we estimated the lagged effects of temperature and precipitation on sporocarp formation and identified the conditions associated with peak fruiting. Although sporocarps formed across a broad range of conditions, peak fruiting was associated with a mean temperature of approximately 13°C averaged over the preceding 20 days and increased linearly with precipitation accumulated over a 26-day window. Under projected warmer and drier autumn conditions, *B. edulis* sporocarp formation in European beech forests is likely to decline.

## Introduction

*Boletus edulis* Bull., commonly known as the porcini, cèpe de Bordeaux, penny bun or Steinpilz, is a Basidiomycete that produces one of the world’s most highly prized edible mushrooms. In parts of Europe, its economic value can even exceed that of timber, contributing substantially to local economic growth (Oria-de-Rueda et al., 2008). Ecologically, *B. edulis* forms ectomycorrhizal associations with a wide variety of host plants, from dominant European coniferous and deciduous tree species to Mediterranean and alpine shrubs (Alonso Ponce et al., 2011; Treindl and Leuchtmann, 2019). In central Europe, it is commonly associated with European beech (*Fagus sylvatica*), the most widespread deciduous tree on the continent, which is increasingly threatened by drought, heat stress and other consequences of climate change (Diers et al., 2023; Leuschner et al., 2023; Martinez del Castillo et al., 2022; Obladen et al., 2021). Ectomycorrhizal fungi (EMF) such as *B. edulis* are thought to bolster tree resilience under such conditions by enhancing water and nutrient uptake, making a better understanding of the biology of EMF an urgent priority in the context of ongoing climate change. Owing to its economic and ecological significance, together with the relative ease of locating its fruiting bodies, *B. edulis* is emerging as a model system for studying EMF (Brejon Lamartinière et al., 2024; Hoffman et al., 2020; Santolamazza-Carbone et al., 2023; Tremble et al., 2023b, 2023a, 2020).

Despite advances in molecular genomics that have illuminated the chromosomal architecture, population structure and evolutionary history of *B. edulis* (Brejon Lamartinière et al., 2025, 2024; Tremble et al., 2023b, 2023a), fundamental aspects of its biology remain poorly understood. This gap partly arises due to our inability to cultivate this species through its complete life cycle under laboratory conditions, which precludes experimental manipulation. In particular, the environmental conditions that trigger sporocarp formation remain elusive, in contrast to many saprotrophic and cultivated fungi, where fruiting conditions have been characterised with remarkable precision under laboratory conditions (Guo and Xu, 2025). For example, the optimal fruiting temperatures for producing multiple *Pleurotus* (oyster mushroom) species have been determined to within a single degree Celsius (Guo and Xu, 2025).

In EMF, where fruiting has not yet been successfully induced in the laboratory, long-term field monitoring of sporocarps has identified temperature and precipitation as the primary drivers of fruiting (Andrew, 2025; Karavani et al., 2018; Martínez-Peña et al., 2012b, 2012a; Tsunoda et al., 2025). However, these variables are not independent and may act jointly (Karavani et al., 2018), requiring their joint inclusion in statistical models to disentangle their individual effects. Moreover, because sporocarp development can take several days (Yang et al., 2025), the environmental conditions that initiate fruiting are likely to act during the period preceding sporocarp emergence and observation. Consequently, modelling sporocarp production requires accounting for the lagged effects of both temperature and precipitation, while determining the length of these potentially distinct lag periods from daily-resolution environmental data and sporocarp observations. To our knowledge, however, no study has modelled EMF sporocarp production at daily resolution while jointly estimating distinct, separately optimised lag periods for the effects of both temperature and precipitation.

In Bielefeld, north-west Germany, a population of *B. edulis* has been continuously monitored since 2015 (Hoffman et al., 2020). The study area, located west of the city, comprises multiple close woodland patches dominated by *F. sylvatica*. Within these patches, sporocarp production has been continuously tracked across 15–29 study sites (see methods for details), with every sporocarp sighting systematically recorded. Using this long-term dataset, we investigated the meteorological conditions associated with sporocarp formation in *B. edulis*. Specifically, we first jointly estimated the optimal lag periods for temperature and precipitation that best predict sporocarp formation using generalised mixed effect models (GLMMs). We then used the model with the best fitting lag periods to estimate the optimal meteorological conditions leading to *B. edulis* sporocarp formation.

## Materials and Methods

### Study population and fieldwork

From 2015 to 2024 inclusive, we monitored *B. edulis* sporocarp production across our study area near Bielefeld (52°03’42.2’N 8°27’03.3’E). The area comprises multiple woodland patches dominated by *F. sylvatic*a followed by common oak (*Quercus robur*) and birch (*Betula pendula*). Monitoring initially focused on 15 study sites, with this number gradually increasing to 29 as additional fruiting locations were discovered, although some earlier sites have since ceased fruiting. Site area ranged from 30 m ^2^ to approximately 1.2 ha (mean = 2,340 m^2^, s.d. = 4,295 m^2^). Sporocarp production was monitored at every active site, with visits conducted 2–4 times a week from early August until the first heavy frost or, alternatively, until approximately two weeks after the last sporocarp was sighted, whichever occurred first (typically around mid-October to mid-November). During each visit, the date and location of every sporocarp was recorded using a handheld GPS device.

### Environmental data

To investigate the meteorological conditions that trigger sporocarp formation, we retrieved the 24h mean temperature (C) and precipitation (mm) for our study area using Copernicus E-OBS *in situ* meteorological data observations at the maximum resolution of 0.1 degrees (Cornes et al., 2018). As all of our study sites fitted within a single grid cell, we extracted the environmental data corresponding to the centroid of the study area using bilinear interpolation, which computes a distance-weighted average from the four nearest cell centres to better approximate local conditions. For subsequent analyses, we considered the period between August 5th–the date of our earliest sporocarp sighting–and November 21st, the date of our latest sporocarp sighting. All downstream analyses were conducted using sporocarp data pooled from all of the sites to maximise statistical power. We additionally repeated the analyses separately for the two most productive sites to explore inter-site variation while retaining adequate statistical power.

### Data analyses

### 1. Lag periods of temperature and precipitation effects

We first conducted an exploratory analysis to identify the appropriate lag periods for the effects of temperature and precipitation on sporocarp formation across the study period. Daily mean temperature and precipitation values were averaged over right-aligned sliding windows of lengths 2, 5, 8, 11, 14, 17, 20, 23, 26, 29, 32 and 35 days. To determine the most appropriate lag periods for both variables, we constructed separate GLMMs predicting daily sporocarp counts as a function of all possible combinations of temperature and precipitation window lengths. These models were specified with a negative binomial error distribution to account for overdispersion and the large number of zero observations (sporocarps were not sighted on 85.8% of days), and included year as a random effect to account for inter-annual variation. Temperature was modelled as a quadratic term to allow for a non-linear relationship with sporocarp production as documented in saprotrophic fungi (Guo and Xu, 2025), while precipitation was modelled as a linear predictor because no upper threshold has yet been identified beyond which additional precipitation no longer promotes sporocarp formation (Tsunoda et al., 2025). The optimal lag periods for temperature and precipitation were then identified by selecting their corresponding window lengths in the best performing model based on Akaike’s Information Criterion (AIC). These models were implemented using the GLMMTMB package (version 1.1.13, Brooks et al., 2017).

### 2. Optimal conditions triggering sporocarp formation

Using the mean temperature and precipitation values computed over the optimal window lengths identified above, we performed two-dimensional kernel density estimation to compare the meteorological conditions associated with sporocarp formation against the full range of meteorological conditions observed during throughout the study period. This comparison allowed us to investigate whether sporocarp occurrence reflects specific climatic preferences, or whether fruiting simply occurs in proportion to the frequency of the prevailing meteorological conditions, such that sporocarps are more common under frequently occurring conditions rather than under biologically preferred ones. Kernel density estimates were computed using the MASS package (version 7.3-65, Venables and Ripley, 2002) and density surfaces were visualised with GGPLOT2 (version 3.5.2, Wickham 2016). Finally, we formally derived the optimal meteorological conditions for sporocarp formation using the best-fitting GLMM described above. All analyses and visualisations were implemented in R v4.5.1 (R core team 2025). To maximise transparency and reproducibility, the data and code used to analyze them have been archived at zenodo (https://zenodo.org/records/20591123).

## Results and discussion

Across the full dataset, the interval between the earliest sporocarp sighting recorded across all years and the latest sporocarp sighting across all years was 86 days. During this overall fruiting window, 24h mean temperatures ranged from -2.5 to 25.9 °C (FIG 1A), while daily mean precipitation varied between zero and 42.6 mm (FIG. 1B). The total number of sporocarps varied substantially among years, ranging from four in 2016 to 354 in 2020 (FIG 1). In several years, fruiting was not continuous but occurred in multiple distinct pulses within a season (e.g. 2020; Fig. 1).

**Figure. 1.**
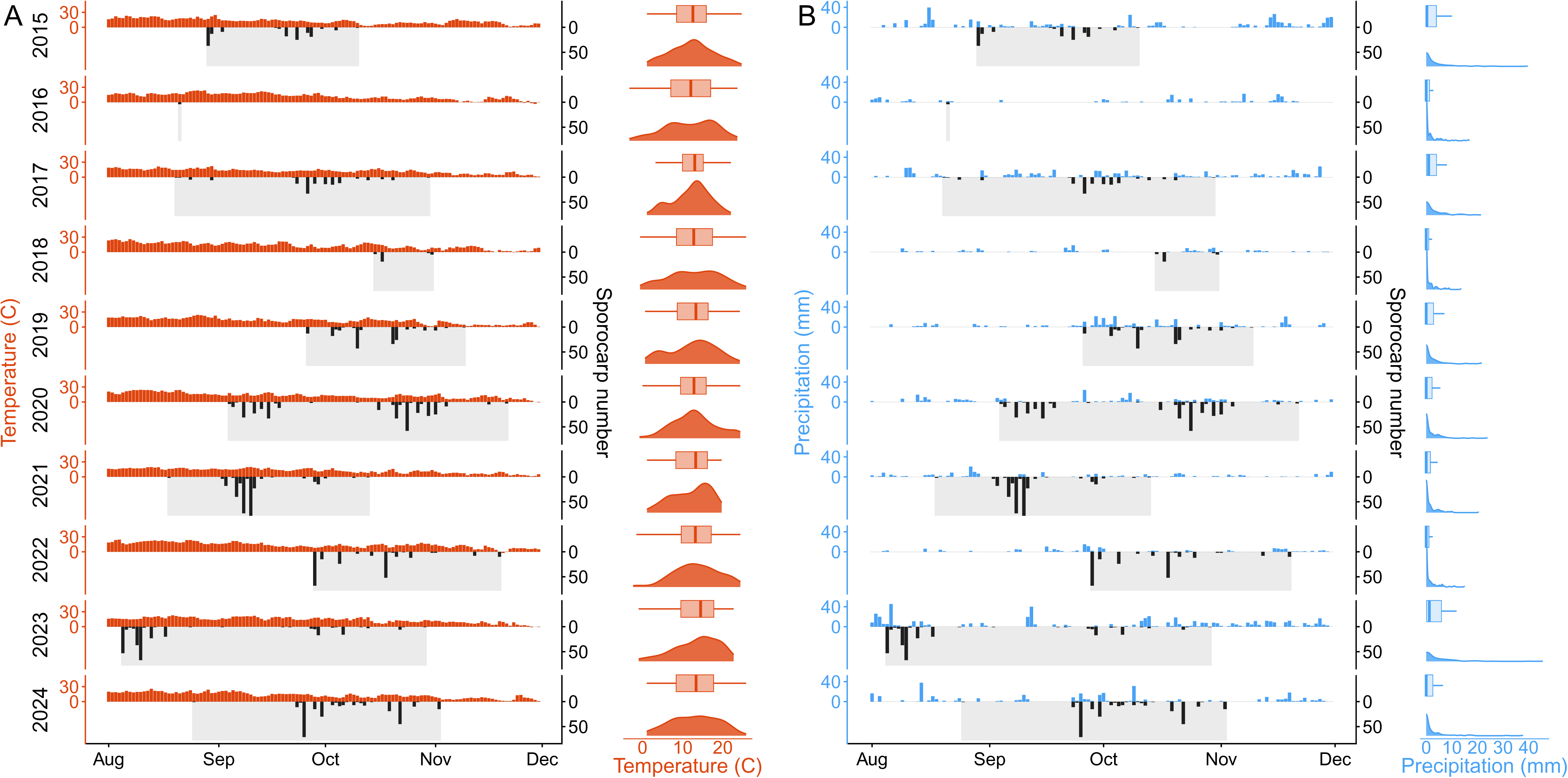
Overview of within- and among-year variation in meteorological conditions and sporocarp observations across the ten-year study period. Panel A shows 24h mean temperature (°C) values above the x axis (in red), while panel (B) shows 24h mean precipitation (mm) values above the x axis (in blue). Daily sporocarp sightings are plotted below the x-axes of both panels (in black). On the right side of each panel, the annual distribution of 24h mean temperature (in red) and precipitation (in blue) values are represented with using simplified boxplots (top), summarizing the first three quartiles interquartile range (box) and the most extreme values within 1.5 times the interquartile range (whiskers), and density plots (bottom), summarising the frequency of observations through a based on single-dimension kernel density estimation. Grey shading areas highlights indicate the interval between the first and last spororocarp sighting of each season. In 2016, sporocarps were only sighted on one day.

Sites were visited frequently (2–4 times per week) and locations known from previous observations to produce sporocarps were consistently revisited, minimising the chance of missed fruiting events. This intensive, targeted monitoring approach made the dataset as exhaustive as practically achievable. The near absence of sporocarps in 2016 therefore appears to be a genuine biological signal rather than a sampling artefact, likely reflecting exceptionally dry conditions during the peak fruiting period (FIG 1B). This interpretation is further supported by the occurrence of moderate fruiting in the preceding and following years (FIG 1). Although total sporocarp counts were generally higher in the latter half of the study, partly because new fruiting sites were discovered over time, no consistent trends in temperature or precipitation were observed across the years. We therefore interpret variation in sporocarp numbers among years as primarily reflecting true biological variation rather than sampling artifacts.

To determine the most appropriate lag periods for the effects of temperature and precipitation on sporocarp production, we constructed separate GLMMs predicting daily sporocarp counts as a function of all possible combinations of window length for both variables (see Methods for details). The best fitting model identified an optimal lag period of 20 days for temperature, both when all sites were analysed jointly, as well as when the two most productive sites were modelled separately (Table S1, Fig. 2, Fig. 3). For precipitation, the best fitting model identified a lag period of 26 days when sites were combined, and 26-32 days when the two largest sites were analysed separately (Table S1, Fig. 2, Fig. 3). These findings indicate that sporocarp formation is triggered most strongly by approximately three weeks of favourable temperature conditions, while the effect of precipitation has a longer lag period, likely reflecting the time needed to build and up and maintain elevated soil moisture levels (Karavani et al., 2018). In line with this, previous studies incorporating both variables similarly reported precipitation lags of approximately one month across multiple EMF species (Karavani et al., 2018), whereas studies excluding temperature yielded shorter estimated lags of less than two weeks for the effects of precipitation (Tsunoda et al., 2025).

**Figure 2.**
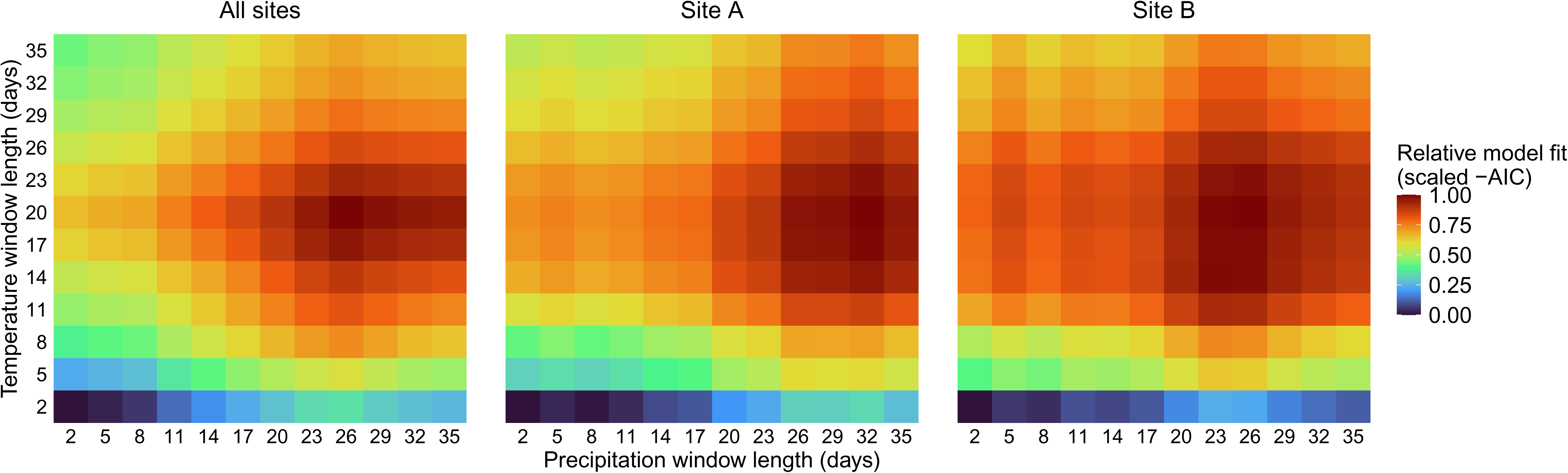
Heatmap of the model fit sensitivity to the sliding window lengths (lag time) used to summarise temperature (y axis) and precipitation (x axis) preceding sporocarp observations. The three panels represent, left to right, the models fitted with the data from all sites combined, the largest site (site A), and the second largest site (site B). The fill represents the Akaike Information Criterion (AIC), scaled to be comparable among the three panels despite sample sizes differences and reversed such that darker red indicates better relative fit.

**Figure. 3.**
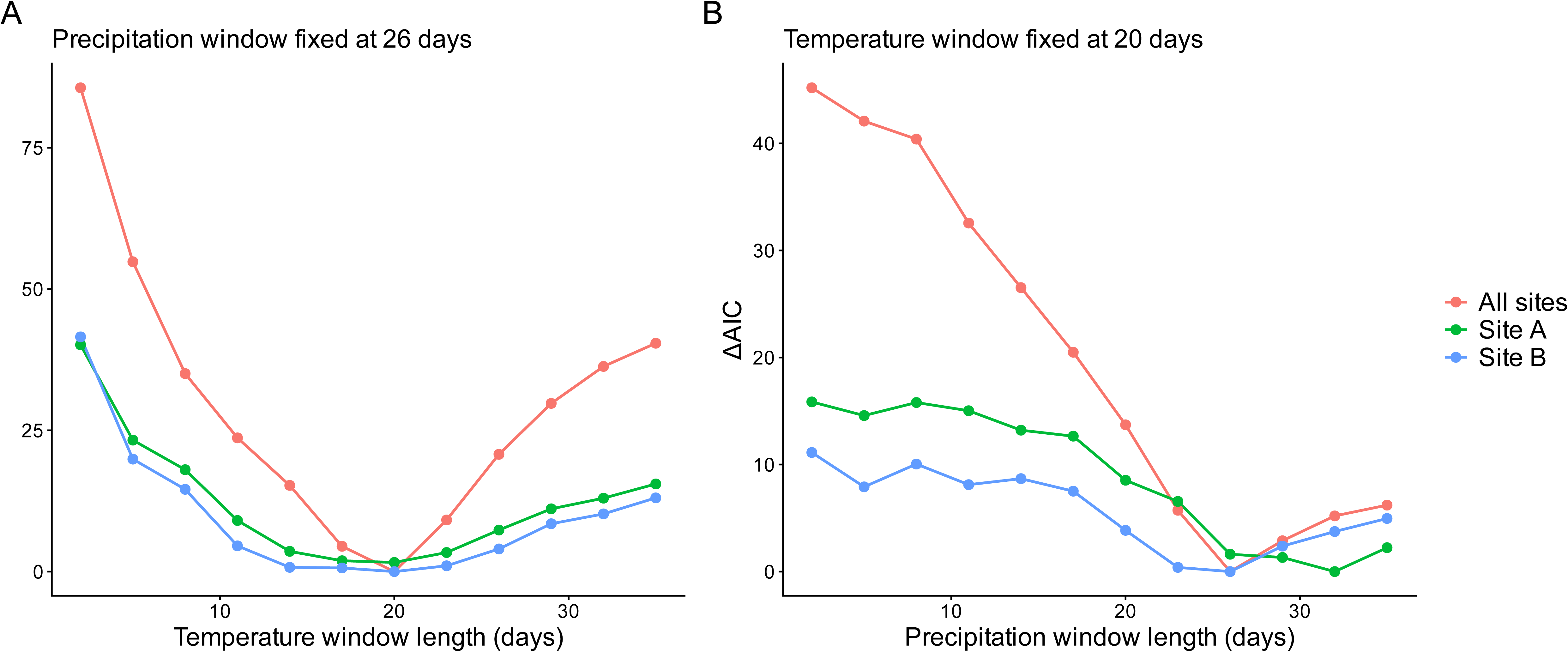
ΔAIC values across sliding-window lengths when the complementary window is held constant at optimum. A) Precipitation window fixed at 26 days, with the temperature window length varying from 2 to 35 days (x axis). B) Temperature window length fixed at 20 days, with the precipitation window length varying from 2 to 35 days (x axis). ΔAIC represents the difference in AIC between each model and the best fitting model for that grouping, such that lower ΔAIC values indicate better model fit. The lines represent the models fitted with the data from all sites combined in red, the most productive site in green and the second most productive site in blue.

To evaluate the meteorological conditions triggering sporocarp formation, we applied two-dimensional kernel density estimation to mean temperature and precipitation values computed over right-aligned sliding windows corresponding to the previously identified optimal lag periods. This analysis was first conducted across all days of the study period to characterise the full range of environmental conditions encountered during monitoring. It was then repeated only using days on which sporocarps were observed to distinguish whether the occurrence of sporocarps reflects specific climatic preferences or whether fruiting simply occurs in proportion to the frequency of the prevailing meteorological conditions. Across the full study period, meteorological conditions spanned a broad range of temperatures (20 day means: 5–20 °C) while precipitation was more constrained, with mean daily values over 35-day windows rarely falling below 1mm or exceeding 4mm (Fig. 4A).

**Figure. 4.**
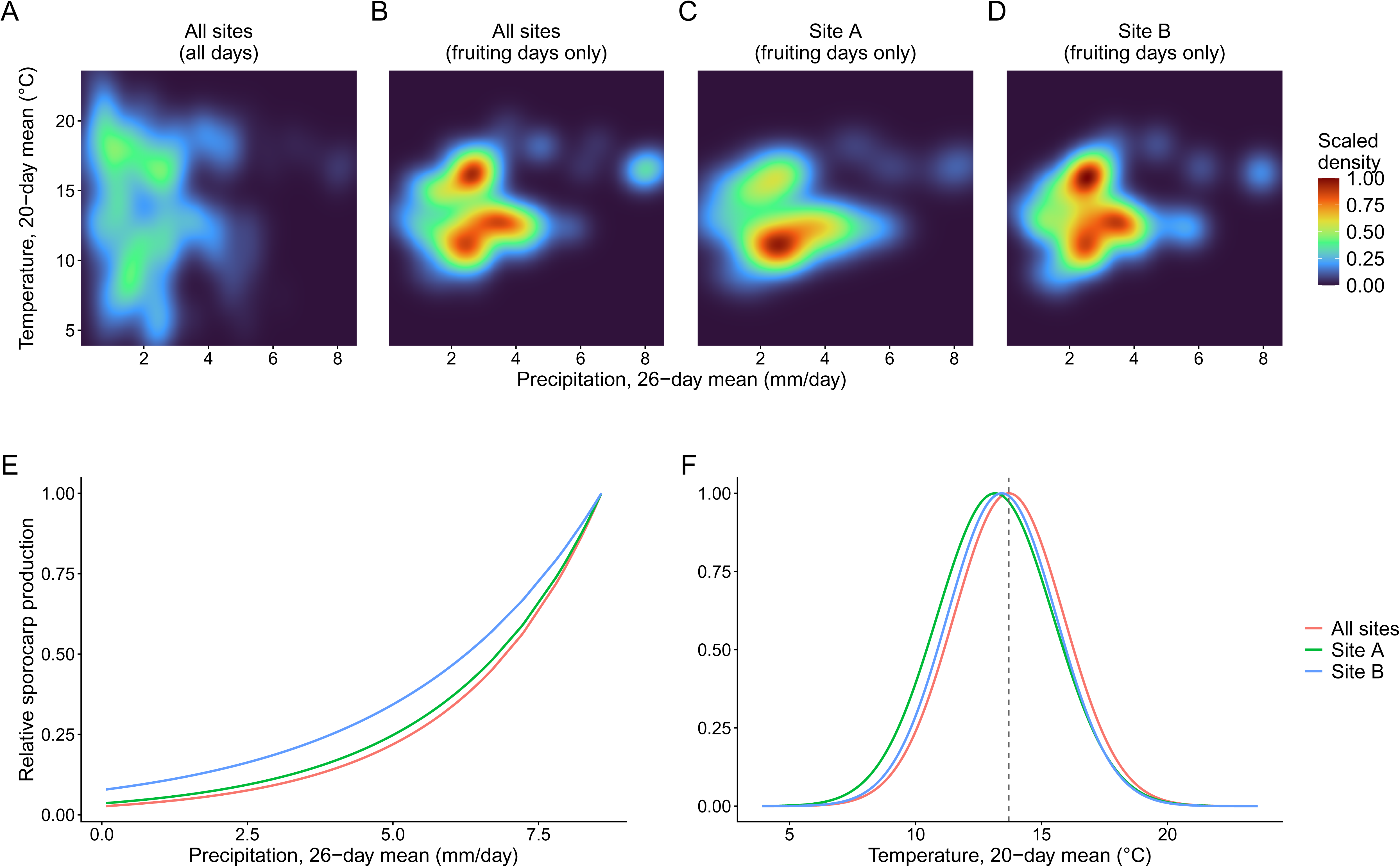
<u>First row</u>: two-dimensional kernel density estimates of the 20-day mean temperature and 26-day mean precipitation of all days of the monitored period A), fruiting days only across all sites combined B), fruiting days only at the most productive site C),and fruiting days only at the second most productive site D). Densities were rescaled to between 0 and 1 within each panel to simplify cross-panel comparison. <u>Second row</u>: Predicted relative sporocarp production as a function of E) the 26-day mean precipitation with temperature held constant at the study mean, and F) the 20-day mean temperature with precipitation held constant at the study mean from the negative binomial generalised linear mixed effect model. The lines represent the models fitted with the data from all sites combined in red, the most productive site in green and the second most productive site in blue. The dashed vertical line represents the optimal temperature across all sites, 13.7 C.

By contrast, fruiting events were concentrated within a narrower range of conditions, particularly temperatures between 10 and 15 °C and precipitation levels between 2 and 4mm per day. Furthermore, fruiting was largely absent under the temperature conditions most frequently observed during the study period (5–10°C). This pattern was consistent both when all sites were analysed together and when the two most productive sites were analysed separately. Notably, two apparent temperature optima were associated with higher sporocarp production (Fig. 4B), a pattern that was also present when sites A and B were analysed separately (Fig. 4C, D). However, these apparent peaks were separated by intermediate temperature and precipitations that were rarely observed during the study period (Fig. 4A), potentially obscuring a continuous underlying response surface. Beyond these apparent peaks, there was no evidence that the effect of temperature depended on precipitation or *vice versa*, arguing against the inclusion of an interaction term in the subsequent models.

To formally quantify relationships between the meteorological variables and sporocarp formation while accounting for repeated observations across years and sites, we used the best fitting GLMMs: one including all sites and two additional models fitted respectively for the two most productive sites. Precipitation had a consistently positive effect on sporocarp formation across all models, in agreement with the results of the two-dimensional kernel density estimation and the general expectation that higher rainfall should promote sporocarp formation (Fig. 4B4E). This finding is also consistent with previous studies of other EMF, which likewise found no evidence for an upper precipitation threshold beyond which fruiting declines (Tsunoda et al., 2025).

By contrast, temperature exhibited a significant quadratic relationship with sporocarp production, revealing an optimum at approximately 13 °C (Fig. 4F) that differed by less than 0.6 °C among the three models (Table S1). This predicted optimum is similar to the temperature used to trigger sporocarp formation in multiple species of cultivated saprotrophic fungi including *Pleurotus*, *Morchella* and *Hypsizygus* spp. (Guo and Xu, 2025). Because these taxa span distantly related fungal families with distinct ecologies, this similarity suggests that the regulatory pathways governing temperature-dependent sporocarp formation may be broadly conserved across mushroom-forming fungi. This convergence is perhaps not unexpected given morphological similarities among the sporocarps of mycorrhizal and saprotrophic basidiomycetes.

## Conclusion

In this study, we identified the meteorological conditions most strongly associated with *B. edulis* sporocarp formation in a beech woodland near Bielefeld, Germany. Temperature emerged as the main short-term predictor, with a 20-day mean temperature of approximately 13°C being associated with the highest probability of sporocarp formation. By contrast, the effect of precipitation was not immediate but accumulated over a longer period, with elevated rainfall over the preceding 26 days increasing the likelihood of fruiting. In the context of environmental change, meteorological conditions in many regions are expected to shift away from the temperature ranges currently associated with optimal sporocarp production in many saprotrophic fungi (Guo and Xu, 2025). Our results suggest that a similar change may also occur for *B. edulis* populations across much of central Europe. However, occasional sightings of sporocarps at higher temperatures in our study suggest that some capacity for persistence may exist, whether mediated by phenotypic plasticity or evolutionary adaptation. Long-term monitoring across broader climatic gradients will be required to determine the extent to which such responses can buffer future declines in fruiting.

## Acknowledgements

We thank Vincent Bouyer for his support in providing access to woodland areas for this study.

## Data availability statement

All the data and code used to produce the analysis and the figures have been archived at Zotero (https://zenodo.org/records/20591123) and Github (https://github.com/ebrejonl/Boletus_fruiting_triggers).

## Supplementary for the manuscript

**Table S1:** Sensitivity of model fit and parameter estimates to the temperature and precipitation window lengths. Each row is one combination of temperature (win_T) and precipitation (win_P) right-aligned sliding-window lengths (days), used to summarise the meteorological conditions preceding daily sporocarp counts. For each combination, a separate model was fitted with the data from all sites and from the two most productive sites (site A, site B) separately. The columns AIC and dAIC represent the Akaike’s Information Criterion and the difference from the best-fitting model within each site (best model = 0). The columns coef_P, se_P and pval_P give the estimate, standard error and p-value for the effect of precipitation. Since Temperature was fitted as a quadratic term, it comprises a linear term (T1) and a quadratic term (T2). Therefore, columns coef_T1, se_T1, pval_T1 and coef_T2, se_T2, pval_T2 give the estimate, standard error and p-value of the linear term and the quadratic term, respectively. Finally, peak_T represents the model-predicted optimal temperature in Celsius. All values are rounded to five decimals.

| win T | win P | AIC | coef P | se P | pval P | coef T1 | se T1 | pval T1 | coef T2 | se T2 | pval T2 | peak T | site | dAIC |
| --- | --- | --- | --- | --- | --- | --- | --- | --- | --- | --- | --- | --- | --- | --- |
| 2 | 2 | 1904.82865 | 0.02179 | 0.01893 | 0.2497 | 0.68017 | 0.11816 | 0 | -0.02649 | 0.00452 | 0 | 12.84049 | All | 131.99089 |
| 5 | 2 | 1874.23154 | 0.03061 | 0.0187 | 0.1016 | 1.13217 | 0.16639 | 0 | -0.0431 | 0.00623 | 0 | 13.13329 | All | 101.39378 |
| 8 | 2 | 1854.06773 | 0.03255 | 0.01873 | 0.0823 | 1.47431 | 0.20388 | 0 | -0.05616 | 0.00759 | 0 | 13.12626 | All | 81.22997 |
| 11 | 2 | 1842.71958 | 0.02944 | 0.01897 | 0.12068 | 1.71976 | 0.23045 | 0 | -0.06531 | 0.00852 | 0 | 13.16655 | All | 69.88182 |
| 14 | 2 | 1833.41199 | 0.02647 | 0.01895 | 0.16249 | 1.97131 | 0.25553 | 0 | -0.07431 | 0.00937 | 0 | 13.26494 | All | 60.57423 |
| 17 | 2 | 1822.97558 | 0.02341 | 0.0189 | 0.21549 | 2.29856 | 0.29057 | 0 | -0.08539 | 0.0105 | 0 | 13.4593 | All | 50.13782 |
| 20 | 2 | 1818.03329 | 0.02057 | 0.01901 | 0.27913 | 2.58993 | 0.32221 | 0 | -0.0947 | 0.01147 | 0 | 13.67377 | All | 45.19552 |
| 23 | 2 | 1823.81716 | 0.02066 | 0.01904 | 0.27803 | 2.65613 | 0.32934 | 0 | -0.09618 | 0.01162 | 0 | 13.80845 | All | 50.97939 |
| 26 | 2 | 1832.18741 | 0.02298 | 0.01914 | 0.22981 | 2.65942 | 0.32983 | 0 | -0.09536 | 0.01154 | 0 | 13.94447 | All | 59.34965 |
| 29 | 2 | 1839.81994 | 0.02504 | 0.01921 | 0.19239 | 2.6882 | 0.33367 | 0 | -0.09536 | 0.01157 | 0 | 14.09444 | All | 66.98218 |
| 32 | 2 | 1845.36726 | 0.02578 | 0.01925 | 0.18047 | 2.73618 | 0.33922 | 0 | -0.09637 | 0.01168 | 0 | 14.19686 | All | 72.5295 |
| 35 | 2 | 1848.60829 | 0.02595 | 0.0192 | 0.17639 | 2.82717 | 0.34952 | 0 | -0.09891 | 0.01196 | 0 | 14.2915 | All | 75.77053 |
| 2 | 5 | 1901.58856 | 0.05918 | 0.02677 | 0.02704 | 0.66572 | 0.11812 | 0 | -0.02584 | 0.00452 | 0 | 12.88202 | All | 128.75079 |
| 5 | 5 | 1871.54787 | 0.06331 | 0.02667 | 0.01759 | 1.11634 | 0.16727 | 0 | -0.04244 | 0.00626 | 0 | 13.1529 | All | 98.7101 |
| 8 | 5 | 1850.24842 | 0.07196 | 0.02661 | 0.00684 | 1.45801 | 0.20411 | 0 | -0.05554 | 0.00759 | 0 | 13.12685 | All | 77.41066 |
| 11 | 5 | 1838.61729 | 0.07113 | 0.02693 | 0.00826 | 1.69971 | 0.23007 | 0 | -0.06458 | 0.0085 | 0 | 13.16036 | All | 65.77952 |
| 14 | 5 | 1829.94403 | 0.06505 | 0.02707 | 0.01627 | 1.9408 | 0.25435 | 0 | -0.07318 | 0.00932 | 0 | 13.25971 | All | 57.10626 |
| 17 | 5 | 1819.7114 | 0.06173 | 0.02731 | 0.02377 | 2.26624 | 0.28984 | 0 | -0.0842 | 0.01047 | 0 | 13.45761 | All | 46.87363 |
| 20 | 5 | 1814.91731 | 0.05883 | 0.02754 | 0.03268 | 2.55719 | 0.32193 | 0 | -0.09351 | 0.01146 | 0 | 13.67384 | All | 42.07955 |
| 23 | 5 | 1820.79421 | 0.05895 | 0.02794 | 0.03485 | 2.61929 | 0.32871 | 0 | -0.09485 | 0.0116 | 0 | 13.80792 | All | 47.95644 |
| 26 | 5 | 1828.91128 | 0.06303 | 0.02816 | 0.0252 | 2.61776 | 0.32837 | 0 | -0.09388 | 0.01149 | 0 | 13.9414 | All | 56.07352 |
| 29 | 5 | 1836.1592 | 0.06783 | 0.02842 | 0.017 | 2.64116 | 0.33121 | 0 | -0.09371 | 0.01148 | 0 | 14.09148 | All | 63.32144 |
| 32 | 5 | 1841.44367 | 0.07021 | 0.02847 | 0.01365 | 2.68774 | 0.33619 | 0 | -0.09468 | 0.01158 | 0 | 14.19363 | All | 68.6059 |
| 35 | 5 | 1844.56477 | 0.07117 | 0.02847 | 0.01242 | 2.77587 | 0.34598 | 0 | -0.09716 | 0.01184 | 0 | 14.28483 | All | 71.727 |
| 2 | 8 | 1897.47956 | 0.1016 | 0.03259 | 0.00182 | 0.66924 | 0.11849 | 0 | -0.0259 | 0.00453 | 0 | 12.92146 | All | 124.6418 |
| 5 | 8 | 1868.42979 | 0.09892 | 0.03236 | 0.00224 | 1.12572 | 0.16912 | 0 | -0.04267 | 0.00633 | 0 | 13.19118 | All | 95.59203 |
| 8 | 8 | 1848.52399 | 0.09936 | 0.03242 | 0.00218 | 1.46382 | 0.20643 | 0 | -0.05559 | 0.00767 | 0 | 13.16677 | All | 75.68623 |
| 11 | 8 | 1836.60776 | 0.10066 | 0.03294 | 0.00224 | 1.69911 | 0.23136 | 0 | -0.06442 | 0.00854 | 0 | 13.18856 | All | 63.77 |
| 14 | 8 | 1828.12346 | 0.09273 | 0.03306 | 0.00503 | 1.93202 | 0.25475 | 0 | -0.07275 | 0.00933 | 0 | 13.27831 | All | 55.28569 |
| 17 | 8 | 1818.14999 | 0.08792 | 0.03348 | 0.00863 | 2.25151 | 0.2897 | 0 | -0.08356 | 0.01046 | 0 | 13.47176 | All | 45.31223 |
| 20 | 8 | 1813.2374 | 0.08633 | 0.03389 | 0.01086 | 2.54281 | 0.3219 | 0 | -0.09289 | 0.01146 | 0 | 13.68657 | All | 40.39964 |
| 23 | 8 | 1819.2371 | 0.08602 | 0.03447 | 0.01258 | 2.60266 | 0.32885 | 0 | -0.09416 | 0.0116 | 0 | 13.81992 | All | 46.39933 |
| 26 | 8 | 1827.56716 | 0.08909 | 0.03485 | 0.01057 | 2.5973 | 0.32857 | 0 | -0.09307 | 0.01149 | 0 | 13.95333 | All | 54.72939 |
| 29 | 8 | 1834.82308 | 0.09459 | 0.03522 | 0.00723 | 2.61481 | 0.33106 | 0 | -0.09272 | 0.01147 | 0 | 14.1012 | All | 61.98531 |
| 32 | 8 | 1839.92259 | 0.09932 | 0.03556 | 0.00523 | 2.65477 | 0.33538 | 0 | -0.09346 | 0.01155 | 0 | 14.20318 | All | 67.08483 |
| 35 | 8 | 1842.92732 | 0.10081 | 0.03538 | 0.00438 | 2.73926 | 0.34474 | 0 | -0.09582 | 0.0118 | 0 | 14.29412 | All | 70.08955 |
| 2 | 11 | 1886.93607 | 0.17401 | 0.03705 | 0 | 0.67321 | 0.11914 | 0 | -0.02605 | 0.00455 | 0 | 12.91946 | All | 114.09831 |
| 5 | 11 | 1857.22966 | 0.1726 | 0.03621 | 0 | 1.14454 | 0.17166 | 0 | -0.04333 | 0.00642 | 0 | 13.20814 | All | 84.3919 |
| 8 | 11 | 1838.30991 | 0.1686 | 0.03632 | 0 | 1.4903 | 0.21072 | 0 | -0.05651 | 0.00782 | 0 | 13.18619 | All | 65.47214 |
| 11 | 11 | 1827.9387 | 0.16324 | 0.03708 | 0.00001 | 1.71084 | 0.23476 | 0 | -0.06478 | 0.00866 | 0 | 13.20558 | All | 55.10094 |
| 14 | 11 | 1819.81479 | 0.15492 | 0.03717 | 0.00003 | 1.92657 | 0.25617 | 0 | -0.07248 | 0.00937 | 0 | 13.2904 | All | 46.97702 |
| 17 | 11 | 1810.08794 | 0.15022 | 0.03743 | 0.00006 | 2.24007 | 0.29105 | 0 | -0.08307 | 0.01049 | 0 | 13.48303 | All | 37.25017 |
| 20 | 11 | 1805.3947 | 0.14855 | 0.03777 | 0.00008 | 2.52927 | 0.32331 | 0 | -0.09233 | 0.01149 | 0 | 13.69761 | All | 32.55694 |
| 23 | 11 | 1811.41303 | 0.14978 | 0.03845 | 0.0001 | 2.58133 | 0.32928 | 0 | -0.09336 | 0.01161 | 0 | 13.82521 | All | 38.57527 |
| 26 | 11 | 1819.80331 | 0.15272 | 0.03893 | 0.00009 | 2.56828 | 0.32823 | 0 | -0.09201 | 0.01148 | 0 | 13.95702 | All | 46.96554 |
| 29 | 11 | 1827.10753 | 0.1576 | 0.03941 | 0.00006 | 2.58149 | 0.33032 | 0 | -0.09151 | 0.01145 | 0 | 14.10447 | All | 54.26976 |
| 32 | 11 | 1832.24239 | 0.16209 | 0.03989 | 0.00005 | 2.61491 | 0.33393 | 0 | -0.09205 | 0.0115 | 0 | 14.20306 | All | 59.40463 |
| 35 | 11 | 1835.09705 | 0.16448 | 0.03986 | 0.00004 | 2.69153 | 0.34228 | 0 | -0.09416 | 0.01172 | 0 | 14.29194 | All | 62.25928 |
| 2 | 14 | 1880.65521 | 0.22434 | 0.04182 | 0 | 0.68064 | 0.11978 | 0 | -0.02646 | 0.00458 | 0 | 12.86198 | All | 107.81744 |
| 5 | 14 | 1850.238 | 0.22315 | 0.04054 | 0 | 1.16065 | 0.17285 | 0 | -0.04403 | 0.00646 | 0 | 13.17937 | All | 77.40024 |
| 8 | 14 | 1830.5969 | 0.221 | 0.04019 | 0 | 1.5239 | 0.21437 | 0 | -0.05781 | 0.00795 | 0 | 13.18073 | All | 57.75913 |
| 11 | 14 | 1820.85763 | 0.21404 | 0.04092 | 0 | 1.75356 | 0.24004 | 0 | -0.06637 | 0.00884 | 0 | 13.21084 | All | 48.01987 |
| 14 | 14 | 1813.84242 | 0.20201 | 0.04124 | 0 | 1.95534 | 0.26009 | 0 | -0.0735 | 0.0095 | 0 | 13.30142 | All | 41.00466 |
| 17 | 14 | 1804.02247 | 0.19802 | 0.04143 | 0 | 2.26588 | 0.29469 | 0 | -0.08396 | 0.01061 | 0 | 13.49365 | All | 31.18471 |
| 20 | 14 | 1799.36266 | 0.19622 | 0.04163 | 0 | 2.55943 | 0.32743 | 0 | -0.09335 | 0.01162 | 0 | 13.70887 | All | 26.52489 |
| 23 | 14 | 1805.85284 | 0.19535 | 0.04225 | 0 | 2.60448 | 0.33228 | 0 | -0.09414 | 0.0117 | 0 | 13.83355 | All | 33.01508 |
| 26 | 14 | 1814.50865 | 0.19738 | 0.04277 | 0 | 2.58144 | 0.33015 | 0 | -0.09245 | 0.01153 | 0 | 13.96173 | All | 41.67089 |
| 29 | 14 | 1821.99319 | 0.20152 | 0.04328 | 0 | 2.59298 | 0.33245 | 0 | -0.09189 | 0.01151 | 0 | 14.10925 | All | 49.15542 |
| 32 | 14 | 1827.59728 | 0.20389 | 0.04381 | 0 | 2.6208 | 0.33604 | 0 | -0.09224 | 0.01156 | 0 | 14.20652 | All | 54.75951 |
| 35 | 14 | 1830.77049 | 0.20546 | 0.04403 | 0 | 2.68926 | 0.34385 | 0 | -0.09407 | 0.01176 | 0 | 14.29337 | All | 57.93273 |
| 2 | 17 | 1873.8011 | 0.27289 | 0.04487 | 0 | 0.69178 | 0.12098 | 0 | -0.02701 | 0.00462 | 0 | 12.80728 | All | 100.96334 |
| 5 | 17 | 1843.62897 | 0.26972 | 0.04369 | 0 | 1.17357 | 0.17384 | 0 | -0.04462 | 0.00649 | 0 | 13.1498 | All | 70.79121 |
| 8 | 17 | 1823.83039 | 0.26667 | 0.04321 | 0 | 1.54223 | 0.21642 | 0 | -0.05855 | 0.00801 | 0 | 13.17067 | All | 50.99263 |
| 11 | 17 | 1813.31567 | 0.26363 | 0.04387 | 0 | 1.79987 | 0.24596 | 0 | -0.06809 | 0.00905 | 0 | 13.2167 | All | 40.47791 |
| 14 | 17 | 1806.65856 | 0.25137 | 0.0442 | 0 | 2.01425 | 0.26699 | 0 | -0.07564 | 0.00974 | 0 | 13.3153 | All | 33.8208 |
| 17 | 17 | 1797.78107 | 0.24358 | 0.04446 | 0 | 2.31925 | 0.30153 | 0 | -0.08585 | 0.01085 | 0 | 13.50793 | All | 24.94331 |
| 20 | 17 | 1793.32796 | 0.24024 | 0.0445 | 0 | 2.60604 | 0.33322 | 0 | -0.09497 | 0.01182 | 0 | 13.72061 | All | 20.49019 |
| 23 | 17 | 1800.23281 | 0.23729 | 0.04502 | 0 | 2.64239 | 0.337 | 0 | -0.09544 | 0.01186 | 0 | 13.84386 | All | 27.39505 |
| 26 | 17 | 1809.43955 | 0.23679 | 0.04551 | 0 | 2.61179 | 0.33428 | 0 | -0.09347 | 0.01167 | 0 | 13.97159 | All | 36.60179 |
| 29 | 17 | 1817.0778 | 0.2411 | 0.04617 | 0 | 2.61528 | 0.33609 | 0 | -0.09262 | 0.01162 | 0 | 14.11876 | All | 44.24003 |
| 32 | 17 | 1822.70324 | 0.24448 | 0.04687 | 0 | 2.64197 | 0.33998 | 0 | -0.09291 | 0.01168 | 0 | 14.21763 | All | 49.86548 |
| 35 | 17 | 1826.37033 | 0.24376 | 0.04708 | 0 | 2.7041 | 0.34792 | 0 | -0.09452 | 0.01188 | 0 | 14.30409 | All | 53.53257 |
| 2 | 20 | 1867.50518 | 0.33212 | 0.05064 | 0 | 0.68771 | 0.12085 | 0 | -0.02706 | 0.00462 | 0 | 12.7085 | All | 94.66742 |
| 5 | 20 | 1837.10699 | 0.32706 | 0.04899 | 0 | 1.17776 | 0.17471 | 0 | -0.04497 | 0.00652 | 0 | 13.09455 | All | 64.26922 |
| 8 | 20 | 1817.19105 | 0.32226 | 0.04822 | 0 | 1.55388 | 0.21791 | 0 | -0.05911 | 0.00806 | 0 | 13.14333 | All | 44.35329 |
| 11 | 20 | 1806.18061 | 0.32108 | 0.04888 | 0 | 1.82828 | 0.24936 | 0 | -0.06921 | 0.00916 | 0 | 13.20775 | All | 33.34284 |
| 14 | 20 | 1798.86899 | 0.31176 | 0.04916 | 0 | 2.07231 | 0.27393 | 0 | -0.07779 | 0.00998 | 0 | 13.31989 | All | 26.03123 |
| 17 | 20 | 1789.82211 | 0.30559 | 0.04946 | 0 | 2.40374 | 0.31112 | 0 | -0.08892 | 0.01119 | 0 | 13.51576 | All | 16.98434 |
| 20 | 20 | 1786.54851 | 0.29651 | 0.04945 | 0 | 2.6773 | 0.34165 | 0 | -0.09752 | 0.01212 | 0 | 13.72686 | All | 13.71074 |
| 23 | 20 | 1794.04328 | 0.29032 | 0.04981 | 0 | 2.6905 | 0.34321 | 0 | -0.09714 | 0.01207 | 0 | 13.84823 | All | 21.20552 |
| 26 | 20 | 1803.67182 | 0.28759 | 0.05024 | 0 | 2.64807 | 0.33977 | 0 | -0.09475 | 0.01185 | 0 | 13.97454 | All | 30.83406 |
| 29 | 20 | 1811.75176 | 0.29009 | 0.05092 | 0 | 2.64515 | 0.34159 | 0 | -0.09366 | 0.0118 | 0 | 14.12178 | All | 38.914 |
| 32 | 20 | 1817.57836 | 0.29356 | 0.05177 | 0 | 2.66599 | 0.34545 | 0 | -0.09373 | 0.01185 | 0 | 14.22179 | All | 44.7406 |
| 35 | 20 | 1821.33759 | 0.29329 | 0.05212 | 0 | 2.72721 | 0.35385 | 0 | -0.09529 | 0.01207 | 0 | 14.30974 | All | 48.49983 |
| 2 | 23 | 1860.75953 | 0.39998 | 0.05761 | 0 | 0.69224 | 0.12114 | 0 | -0.02742 | 0.00461 | 0 | 12.62218 | All | 87.92176 |
| 5 | 23 | 1830.95947 | 0.38973 | 0.0556 | 0 | 1.17718 | 0.1744 | 0 | -0.04515 | 0.0065 | 0 | 13.03737 | All | 58.1217 |
| 8 | 23 | 1811.14859 | 0.38068 | 0.05421 | 0 | 1.55474 | 0.21841 | 0 | -0.05935 | 0.00808 | 0 | 13.09837 | All | 38.31083 |
| 11 | 23 | 1799.97129 | 0.37838 | 0.05457 | 0 | 1.84013 | 0.2511 | 0 | -0.06983 | 0.00923 | 0 | 13.17628 | All | 27.13352 |
| 14 | 23 | 1791.97675 | 0.37191 | 0.05491 | 0 | 2.10456 | 0.27774 | 0 | -0.07911 | 0.01012 | 0 | 13.3008 | All | 19.13899 |
| 17 | 23 | 1781.71217 | 0.37177 | 0.0553 | 0 | 2.48319 | 0.3197 | 0 | -0.09193 | 0.0115 | 0 | 13.50602 | All | 8.87441 |
| 20 | 23 | 1778.56611 | 0.36177 | 0.05511 | 0 | 2.77868 | 0.35169 | 0 | -0.10127 | 0.01249 | 0 | 13.71886 | All | 5.72835 |
| 23 | 23 | 1787.44024 | 0.34847 | 0.05525 | 0 | 2.76855 | 0.35161 | 0 | -0.1 | 0.01237 | 0 | 13.84296 | All | 14.60247 |
| 26 | 23 | 1797.74428 | 0.3413 | 0.05538 | 0 | 2.70429 | 0.34628 | 0 | -0.0968 | 0.01207 | 0 | 13.96901 | All | 24.90651 |
| 29 | 23 | 1806.23103 | 0.34137 | 0.05593 | 0 | 2.69393 | 0.34791 | 0 | -0.09543 | 0.01201 | 0 | 14.11537 | All | 33.39327 |
| 32 | 23 | 1812.42263 | 0.34361 | 0.05681 | 0 | 2.71243 | 0.35222 | 0 | -0.0954 | 0.01208 | 0 | 14.21589 | All | 39.58486 |
| 35 | 23 | 1816.24603 | 0.34475 | 0.05748 | 0 | 2.77348 | 0.36112 | 0 | -0.09693 | 0.01231 | 0 | 14.306 | All | 43.40827 |
| 2 | 26 | 1858.45171 | 0.4512 | 0.06483 | 0 | 0.69228 | 0.12136 | 0 | -0.02752 | 0.00461 | 0 | 12.57865 | All | 85.61395 |
| 5 | 26 | 1827.6633 | 0.44864 | 0.06327 | 0 | 1.17934 | 0.17398 | 0 | -0.04531 | 0.00646 | 0 | 13.01446 | All | 54.82554 |
| 8 | 26 | 1807.87944 | 0.43669 | 0.06152 | 0 | 1.55437 | 0.21796 | 0 | -0.05945 | 0.00805 | 0 | 13.07243 | All | 35.04168 |
| 11 | 26 | 1796.50261 | 0.43185 | 0.06127 | 0 | 1.84514 | 0.25174 | 0 | -0.0702 | 0.00925 | 0 | 13.14259 | All | 23.66484 |
| 14 | 26 | 1788.09271 | 0.42556 | 0.06139 | 0 | 2.12174 | 0.27959 | 0 | -0.07994 | 0.0102 | 0 | 13.27136 | All | 15.25494 |
| 17 | 26 | 1777.3128 | 0.4277 | 0.06184 | 0 | 2.52044 | 0.32374 | 0 | -0.09345 | 0.01166 | 0 | 13.48581 | All | 4.47503 |
| 20 | 26 | 1772.83776 | 0.42404 | 0.06175 | 0 | 2.86579 | 0.36005 | 0 | -0.10455 | 0.0128 | 0 | 13.70482 | All | 0 |
| 23 | 26 | 1781.97449 | 0.40804 | 0.06157 | 0 | 2.8646 | 0.36031 | 0 | -0.10356 | 0.01269 | 0 | 13.83077 | All | 9.13673 |
| 26 | 26 | 1793.5908 | 0.39279 | 0.06138 | 0 | 2.77725 | 0.3534 | 0 | -0.09948 | 0.01233 | 0 | 13.95892 | All | 20.75303 |
| 29 | 26 | 1802.60266 | 0.38861 | 0.06159 | 0 | 2.75191 | 0.35377 | 0 | -0.09755 | 0.01222 | 0 | 14.10531 | All | 29.76489 |
| 32 | 26 | 1809.14291 | 0.38876 | 0.0624 | 0 | 2.76573 | 0.358 | 0 | -0.09734 | 0.01228 | 0 | 14.20629 | All | 36.30514 |
| 35 | 26 | 1813.2462 | 0.38891 | 0.06316 | 0 | 2.83099 | 0.36803 | 0 | -0.09899 | 0.01254 | 0 | 14.2993 | All | 40.40844 |
| 2 | 29 | 1864.62421 | 0.46241 | 0.07237 | 0 | 0.67226 | 0.11961 | 0 | -0.02684 | 0.00456 | 0 | 12.52477 | All | 91.78644 |
| 5 | 29 | 1833.96506 | 0.4648 | 0.0713 | 0 | 1.157 | 0.17208 | 0 | -0.04454 | 0.0064 | 0 | 12.9873 | All | 61.1273 |
| 8 | 29 | 1812.60645 | 0.4647 | 0.06991 | 0 | 1.53815 | 0.21562 | 0 | -0.05891 | 0.00796 | 0 | 13.05469 | All | 39.76868 |
| 11 | 29 | 1800.46675 | 0.46134 | 0.06917 | 0 | 1.8383 | 0.25034 | 0 | -0.07002 | 0.00919 | 0 | 13.12643 | All | 27.62899 |
| 14 | 29 | 1791.56759 | 0.45393 | 0.06869 | 0 | 2.12122 | 0.27921 | 0 | -0.08007 | 0.01019 | 0 | 13.24525 | All | 18.72983 |
| 17 | 29 | 1780.75213 | 0.45493 | 0.06884 | 0 | 2.52189 | 0.3237 | 0 | -0.09367 | 0.01166 | 0 | 13.46183 | All | 7.91436 |
| 20 | 29 | 1775.72125 | 0.45391 | 0.06885 | 0 | 2.88665 | 0.36269 | 0 | -0.10544 | 0.0129 | 0 | 13.6882 | All | 2.88349 |
| 23 | 29 | 1783.5931 | 0.4424 | 0.0687 | 0 | 2.92347 | 0.36622 | 0 | -0.10579 | 0.01291 | 0 | 13.81715 | All | 10.75534 |
| 26 | 29 | 1795.27794 | 0.42377 | 0.06822 | 0 | 2.84364 | 0.3594 | 0 | -0.10195 | 0.01255 | 0 | 13.94587 | All | 22.44017 |
| 29 | 29 | 1805.07954 | 0.41269 | 0.0681 | 0 | 2.80692 | 0.35893 | 0 | -0.09957 | 0.01241 | 0 | 14.09495 | All | 32.24178 |
| 32 | 29 | 1811.69489 | 0.41022 | 0.06866 | 0 | 2.8158 | 0.36252 | 0 | -0.09916 | 0.01244 | 0 | 14.19835 | All | 38.85712 |
| 35 | 29 | 1815.91224 | 0.40872 | 0.06942 | 0 | 2.88174 | 0.37264 | 0 | -0.1008 | 0.0127 | 0 | 14.29401 | All | 43.07448 |
| 2 | 32 | 1868.13968 | 0.47679 | 0.07897 | 0 | 0.67185 | 0.11908 | 0 | -0.02684 | 0.00456 | 0 | 12.5143 | All | 95.30192 |
| 5 | 32 | 1839.16469 | 0.47169 | 0.07811 | 0 | 1.13305 | 0.16937 | 0 | -0.04371 | 0.00632 | 0 | 12.96181 | All | 66.32693 |
| 8 | 32 | 1817.71706 | 0.47964 | 0.07791 | 0 | 1.51313 | 0.21249 | 0 | -0.05808 | 0.00787 | 0 | 13.02618 | All | 44.8793 |
| 11 | 32 | 1804.3195 | 0.48727 | 0.07775 | 0 | 1.82567 | 0.24839 | 0 | -0.06965 | 0.00914 | 0 | 13.10568 | All | 31.48174 |
| 14 | 32 | 1794.37192 | 0.48507 | 0.07707 | 0 | 2.13147 | 0.28016 | 0 | -0.08055 | 0.01023 | 0 | 13.23096 | All | 21.53416 |
| 17 | 32 | 1783.16815 | 0.48658 | 0.07677 | 0 | 2.54138 | 0.32566 | 0 | -0.09453 | 0.01175 | 0 | 13.44285 | All | 10.33039 |
| 20 | 32 | 1778.0408 | 0.48504 | 0.07655 | 0 | 2.91234 | 0.36528 | 0 | -0.10652 | 0.01301 | 0 | 13.67058 | All | 5.20304 |
| 23 | 32 | 1785.31438 | 0.4752 | 0.07635 | 0 | 2.96723 | 0.37078 | 0 | -0.10748 | 0.01309 | 0 | 13.80399 | All | 12.47661 |
| 26 | 32 | 1796.23582 | 0.4584 | 0.07569 | 0 | 2.90645 | 0.36526 | 0 | -0.10431 | 0.01278 | 0 | 13.93233 | All | 23.39806 |
| 29 | 32 | 1806.25008 | 0.44358 | 0.07526 | 0 | 2.86906 | 0.36424 | 0 | -0.10187 | 0.01262 | 0 | 14.08146 | All | 33.41232 |
| 32 | 32 | 1813.39745 | 0.43536 | 0.07547 | 0 | 2.86972 | 0.36708 | 0 | -0.10112 | 0.01262 | 0 | 14.18912 | All | 40.55969 |
| 35 | 32 | 1817.6061 | 0.43172 | 0.07592 | 0 | 2.93419 | 0.37693 | 0 | -0.10268 | 0.01286 | 0 | 14.28808 | All | 44.76833 |
| 2 | 35 | 1870.31492 | 0.49326 | 0.08517 | 0 | 0.68041 | 0.11963 | 0 | -0.02718 | 0.00458 | 0 | 12.51732 | All | 97.47715 |
| 5 | 35 | 1840.92217 | 0.49531 | 0.08526 | 0 | 1.14142 | 0.16879 | 0 | -0.04411 | 0.00631 | 0 | 12.93853 | All | 68.08441 |
| 8 | 35 | 1820.08923 | 0.50294 | 0.08572 | 0 | 1.5091 | 0.20972 | 0 | -0.05807 | 0.00779 | 0 | 12.99419 | All | 47.25147 |
| 11 | 35 | 1806.79788 | 0.51583 | 0.08667 | 0 | 1.82195 | 0.2454 | 0 | -0.06967 | 0.00906 | 0 | 13.07539 | All | 33.96011 |
| 14 | 35 | 1796.08105 | 0.52253 | 0.08672 | 0 | 2.13723 | 0.27877 | 0 | -0.08091 | 0.01021 | 0 | 13.20821 | All | 23.24328 |
| 17 | 35 | 1784.21205 | 0.52942 | 0.08649 | 0 | 2.5662 | 0.32622 | 0 | -0.09557 | 0.01179 | 0 | 13.4255 | All | 11.37429 |
| 20 | 35 | 1779.04586 | 0.52703 | 0.08597 | 0 | 2.94386 | 0.36662 | 0 | -0.10783 | 0.01308 | 0 | 13.65001 | All | 6.2081 |
| 23 | 35 | 1786.56389 | 0.51237 | 0.08524 | 0 | 2.99981 | 0.37254 | 0 | -0.1088 | 0.01317 | 0 | 13.78535 | All | 13.72613 |
| 26 | 35 | 1797.00993 | 0.49583 | 0.08423 | 0 | 2.95581 | 0.36877 | 0 | -0.10619 | 0.01292 | 0 | 13.91722 | All | 24.17217 |
| 29 | 35 | 1806.64262 | 0.4807 | 0.08337 | 0 | 2.92951 | 0.36858 | 0 | -0.10416 | 0.01279 | 0 | 14.06285 | All | 33.80486 |
| 32 | 35 | 1814.06744 | 0.46781 | 0.08306 | 0 | 2.92821 | 0.37104 | 0 | -0.10333 | 0.01278 | 0 | 14.16875 | All | 41.22968 |
| 35 | 35 | 1818.62088 | 0.4593 | 0.08304 | 0 | 2.99092 | 0.38077 | 0 | -0.10478 | 0.01302 | 0 | 14.2724 | All | 45.78312 |
| 2 | 2 | 858.33355 | 0.01164 | 0.0339 | 0.73124 | 0.47007 | 0.15609 | 0.0026 | -0.01989 | 0.00622 | 0.00137 | 11.81448 | Site_A | 58.62309 |
| 5 | 2 | 839.81624 | 0.01691 | 0.03349 | 0.61354 | 1.06242 | 0.25549 | 0.00003 | -0.04276 | 0.00993 | 0.00002 | 12.42263 | Site_A | 40.10578 |
| 8 | 2 | 833.79635 | 0.01936 | 0.03353 | 0.56363 | 1.27323 | 0.29262 | 0.00001 | -0.05147 | 0.01136 | 0.00001 | 12.36954 | Site_A | 34.08589 |
| 11 | 2 | 824.2589 | 0.01564 | 0.03397 | 0.64523 | 1.67178 | 0.36555 | 0 | -0.0665 | 0.014 | 0 | 12.56913 | Site_A | 24.54844 |
| 14 | 2 | 818.29706 | 0.01126 | 0.03397 | 0.74033 | 2.01023 | 0.42441 | 0 | -0.07884 | 0.01607 | 0 | 12.74938 | Site_A | 18.5866 |
| 17 | 2 | 816.32044 | 0.008 | 0.03394 | 0.81362 | 2.18892 | 0.45654 | 0 | -0.08493 | 0.01707 | 0 | 12.88613 | Site_A | 16.60997 |
| 20 | 2 | 815.56394 | 0.00479 | 0.03414 | 0.88853 | 2.3759 | 0.49318 | 0 | -0.09077 | 0.01814 | 0 | 13.08803 | Site_A | 15.85348 |
| 23 | 2 | 816.6322 | 0.00387 | 0.03425 | 0.90996 | 2.51037 | 0.51668 | 0 | -0.09446 | 0.01874 | 0 | 13.28826 | Site_A | 16.92174 |
| 26 | 2 | 819.87871 | 0.00505 | 0.03442 | 0.88334 | 2.5344 | 0.51895 | 0 | -0.09416 | 0.01859 | 0 | 13.45777 | Site_A | 20.16825 |
| 29 | 2 | 823.28152 | 0.007 | 0.03448 | 0.8391 | 2.55875 | 0.52117 | 0 | -0.09393 | 0.01847 | 0 | 13.62104 | Site_A | 23.57106 |
| 32 | 2 | 824.92645 | 0.00767 | 0.03441 | 0.82366 | 2.6545 | 0.53372 | 0 | -0.09648 | 0.01875 | 0 | 13.75739 | Site_A | 25.21599 |
| 35 | 2 | 827.10623 | 0.0086 | 0.03431 | 0.80201 | 2.70585 | 0.5415 | 0 | -0.09755 | 0.01887 | 0 | 13.86862 | Site_A | 27.39577 |
| 2 | 5 | 856.45105 | 0.06584 | 0.04457 | 0.13955 | 0.45223 | 0.15536 | 0.0036 | -0.01909 | 0.00619 | 0.00204 | 11.84735 | Site_A | 56.74059 |
| 5 | 5 | 838.2525 | 0.06244 | 0.04451 | 0.16065 | 1.03962 | 0.25616 | 0.00005 | -0.04182 | 0.00995 | 0.00003 | 12.4292 | Site_A | 38.54204 |
| 8 | 5 | 831.76364 | 0.07162 | 0.04455 | 0.10787 | 1.25037 | 0.29231 | 0.00002 | -0.05061 | 0.01135 | 0.00001 | 12.35337 | Site_A | 32.05318 |
| 11 | 5 | 822.22069 | 0.07077 | 0.04527 | 0.11798 | 1.64519 | 0.3648 | 0.00001 | -0.06557 | 0.01398 | 0 | 12.54582 | Site_A | 22.51023 |
| 14 | 5 | 816.61527 | 0.06316 | 0.04544 | 0.16454 | 1.97346 | 0.42268 | 0 | -0.07752 | 0.01602 | 0 | 12.72929 | Site_A | 16.90481 |
| 17 | 5 | 814.85143 | 0.05849 | 0.04572 | 0.20085 | 2.14747 | 0.45525 | 0 | -0.08344 | 0.01704 | 0 | 12.86875 | Site_A | 15.14097 |
| 20 | 5 | 814.28517 | 0.05421 | 0.04607 | 0.23929 | 2.32551 | 0.49145 | 0 | -0.08896 | 0.01809 | 0 | 13.07088 | Site_A | 14.57471 |
| 23 | 5 | 815.46732 | 0.05217 | 0.04667 | 0.26362 | 2.44797 | 0.51379 | 0 | -0.09225 | 0.01865 | 0 | 13.26871 | Site_A | 15.75686 |
| 26 | 5 | 818.63276 | 0.05459 | 0.04701 | 0.24548 | 2.46108 | 0.51407 | 0 | -0.09159 | 0.01843 | 0 | 13.43509 | Site_A | 18.9223 |
| 29 | 5 | 821.8911 | 0.05853 | 0.04736 | 0.2165 | 2.4787 | 0.51515 | 0 | -0.09114 | 0.01826 | 0 | 13.59877 | Site_A | 22.18064 |
| 32 | 5 | 823.43399 | 0.06076 | 0.04734 | 0.19931 | 2.57097 | 0.52683 | 0 | -0.09359 | 0.01852 | 0 | 13.73496 | Site_A | 23.72353 |
| 35 | 5 | 825.55714 | 0.06193 | 0.04717 | 0.18916 | 2.62038 | 0.53426 | 0 | -0.09466 | 0.01864 | 0 | 13.84095 | Site_A | 25.84668 |
| 2 | 8 | 857.7981 | 0.04776 | 0.05775 | 0.40825 | 0.46419 | 0.1561 | 0.00294 | -0.01957 | 0.00622 | 0.00165 | 11.86276 | Site_A | 58.08764 |
| 5 | 8 | 839.64266 | 0.03803 | 0.05768 | 0.5097 | 1.05856 | 0.25693 | 0.00004 | -0.04251 | 0.00998 | 0.00002 | 12.45023 | Site_A | 39.9322 |
| 8 | 8 | 833.70992 | 0.03755 | 0.05802 | 0.51748 | 1.26539 | 0.29371 | 0.00002 | -0.05108 | 0.01139 | 0.00001 | 12.38733 | Site_A | 33.99946 |
| 11 | 8 | 824.07565 | 0.03706 | 0.05853 | 0.52662 | 1.66232 | 0.3663 | 0.00001 | -0.06609 | 0.01403 | 0 | 12.57713 | Site_A | 24.36519 |
| 14 | 8 | 818.16902 | 0.02887 | 0.05878 | 0.62337 | 1.99829 | 0.42487 | 0 | -0.07835 | 0.01609 | 0 | 12.7519 | Site_A | 18.45856 |
| 17 | 8 | 816.23939 | 0.02213 | 0.05951 | 0.70998 | 2.17728 | 0.4574 | 0 | -0.08448 | 0.01711 | 0 | 12.88694 | Site_A | 16.52893 |
| 20 | 8 | 815.5048 | 0.01696 | 0.06003 | 0.77756 | 2.36512 | 0.49434 | 0 | -0.09035 | 0.01819 | 0 | 13.08836 | Site_A | 15.79434 |
| 23 | 8 | 816.60221 | 0.01263 | 0.06077 | 0.83542 | 2.50091 | 0.51871 | 0 | -0.0941 | 0.01882 | 0 | 13.28847 | Site_A | 16.89175 |
| 26 | 8 | 819.85639 | 0.0129 | 0.06143 | 0.83368 | 2.52445 | 0.5222 | 0 | -0.09379 | 0.01872 | 0 | 13.45794 | Site_A | 20.14593 |
| 29 | 8 | 823.25579 | 0.01607 | 0.06199 | 0.79549 | 2.54477 | 0.52557 | 0 | -0.09342 | 0.01863 | 0 | 13.62059 | Site_A | 23.54533 |
| 32 | 8 | 824.88431 | 0.01898 | 0.06246 | 0.76122 | 2.63552 | 0.5384 | 0 | -0.09579 | 0.01892 | 0 | 13.75669 | Site_A | 25.17385 |
| 35 | 8 | 827.04468 | 0.02198 | 0.06207 | 0.72321 | 2.68251 | 0.54589 | 0 | -0.09672 | 0.01903 | 0 | 13.86738 | Site_A | 27.33422 |
| 2 | 11 | 856.04022 | 0.10321 | 0.06408 | 0.10724 | 0.45754 | 0.15617 | 0.00339 | -0.01925 | 0.00621 | 0.00194 | 11.8871 | Site_A | 56.32976 |
| 5 | 11 | 838.07657 | 0.09247 | 0.06329 | 0.144 | 1.05458 | 0.25834 | 0.00004 | -0.04224 | 0.01002 | 0.00002 | 12.4839 | Site_A | 38.3661 |
| 8 | 11 | 832.35256 | 0.08773 | 0.06391 | 0.16985 | 1.2619 | 0.29562 | 0.00002 | -0.05082 | 0.01145 | 0.00001 | 12.41504 | Site_A | 32.6421 |
| 11 | 11 | 822.96834 | 0.08127 | 0.06457 | 0.20817 | 1.65395 | 0.36788 | 0.00001 | -0.06566 | 0.01407 | 0 | 12.59404 | Site_A | 23.25788 |
| 14 | 11 | 817.18491 | 0.07335 | 0.06478 | 0.25751 | 1.97955 | 0.42496 | 0 | -0.07757 | 0.01608 | 0 | 12.75958 | Site_A | 17.47445 |
| 17 | 11 | 815.36923 | 0.06707 | 0.06538 | 0.30496 | 2.15012 | 0.45647 | 0 | -0.08341 | 0.01706 | 0 | 12.88957 | Site_A | 15.65877 |
| 20 | 11 | 814.73652 | 0.06195 | 0.06592 | 0.34733 | 2.33106 | 0.49269 | 0 | -0.08905 | 0.01813 | 0 | 13.08823 | Site_A | 15.02606 |
| 23 | 11 | 815.89312 | 0.05904 | 0.06676 | 0.37651 | 2.45765 | 0.51577 | 0 | -0.0925 | 0.01872 | 0 | 13.28522 | Site_A | 16.18266 |
| 26 | 11 | 819.15288 | 0.05947 | 0.06748 | 0.37814 | 2.47268 | 0.51766 | 0 | -0.0919 | 0.01856 | 0 | 13.45298 | Site_A | 19.44242 |
| 29 | 11 | 822.53669 | 0.06162 | 0.06818 | 0.36607 | 2.48741 | 0.52013 | 0 | -0.09135 | 0.01844 | 0 | 13.61473 | Site_A | 22.82623 |
| 32 | 11 | 824.15144 | 0.06386 | 0.06897 | 0.35449 | 2.57284 | 0.53244 | 0 | -0.09356 | 0.01871 | 0 | 13.7491 | Site_A | 24.44098 |
| 35 | 11 | 826.27438 | 0.06652 | 0.06891 | 0.33443 | 2.61617 | 0.54 | 0 | -0.0944 | 0.01883 | 0 | 13.85751 | Site_A | 26.56392 |
| 2 | 14 | 853.23679 | 0.1661 | 0.06995 | 0.01757 | 0.45353 | 0.15663 | 0.00379 | -0.01912 | 0.00622 | 0.00212 | 11.8594 | Site_A | 53.52633 |
| 5 | 14 | 835.35172 | 0.15478 | 0.06853 | 0.0239 | 1.05634 | 0.26009 | 0.00005 | -0.04228 | 0.01006 | 0.00003 | 12.49096 | Site_A | 35.64126 |
| 8 | 14 | 829.74978 | 0.14981 | 0.06892 | 0.02973 | 1.26655 | 0.29866 | 0.00002 | -0.05093 | 0.01154 | 0.00001 | 12.43501 | Site_A | 30.03932 |
| 11 | 14 | 820.59736 | 0.14225 | 0.06975 | 0.04141 | 1.66401 | 0.37321 | 0.00001 | -0.06594 | 0.01424 | 0 | 12.61839 | Site_A | 20.8869 |
| 14 | 14 | 815.18396 | 0.13031 | 0.07034 | 0.06394 | 1.97698 | 0.42856 | 0 | -0.07736 | 0.01618 | 0 | 12.77806 | Site_A | 15.4735 |
| 17 | 14 | 813.45459 | 0.12483 | 0.07091 | 0.07834 | 2.13594 | 0.45848 | 0 | -0.08279 | 0.01711 | 0 | 12.89997 | Site_A | 13.74413 |
| 20 | 14 | 812.91933 | 0.11963 | 0.07121 | 0.09296 | 2.30979 | 0.49394 | 0 | -0.0882 | 0.01815 | 0 | 13.0946 | Site_A | 13.20887 |
| 23 | 14 | 814.15599 | 0.11663 | 0.07188 | 0.10467 | 2.42867 | 0.51551 | 0 | -0.09139 | 0.01869 | 0 | 13.28768 | Site_A | 14.44553 |
| 26 | 14 | 817.4309 | 0.11726 | 0.07256 | 0.10607 | 2.43416 | 0.51542 | 0 | -0.09047 | 0.01847 | 0 | 13.45221 | Site_A | 17.72044 |
| 29 | 14 | 820.82011 | 0.11916 | 0.07324 | 0.10374 | 2.44223 | 0.51649 | 0 | -0.0897 | 0.0183 | 0 | 13.61333 | Site_A | 21.10965 |
| 32 | 14 | 822.50449 | 0.11987 | 0.07416 | 0.106 | 2.52066 | 0.52777 | 0 | -0.09169 | 0.01853 | 0 | 13.74567 | Site_A | 22.79403 |
| 35 | 14 | 824.68668 | 0.12066 | 0.07447 | 0.10518 | 2.55688 | 0.53475 | 0 | -0.09231 | 0.01864 | 0 | 13.85003 | Site_A | 24.97622 |
| 2 | 17 | 852.19784 | 0.19857 | 0.07632 | 0.00927 | 0.45195 | 0.15684 | 0.00396 | -0.01915 | 0.00622 | 0.00209 | 11.80193 | Site_A | 52.48738 |
| 5 | 17 | 834.48981 | 0.1841 | 0.07493 | 0.014 | 1.05654 | 0.26088 | 0.00005 | -0.04233 | 0.01008 | 0.00003 | 12.4787 | Site_A | 34.77935 |
| 8 | 17 | 828.91628 | 0.178 | 0.0751 | 0.01778 | 1.26913 | 0.30038 | 0.00002 | -0.05103 | 0.01159 | 0.00001 | 12.43547 | Site_A | 29.20581 |
| 11 | 17 | 819.71204 | 0.17171 | 0.07589 | 0.02367 | 1.68072 | 0.3781 | 0.00001 | -0.0665 | 0.01439 | 0 | 12.63691 | Site_A | 20.00158 |
| 14 | 17 | 814.36896 | 0.1593 | 0.07663 | 0.03764 | 2.00509 | 0.43504 | 0 | -0.0783 | 0.0164 | 0 | 12.80344 | Site_A | 14.6585 |
| 17 | 17 | 812.86573 | 0.14943 | 0.07734 | 0.05333 | 2.15704 | 0.4638 | 0 | -0.08347 | 0.01729 | 0 | 12.92171 | Site_A | 13.15526 |
| 20 | 17 | 812.35987 | 0.14318 | 0.0774 | 0.06435 | 2.32557 | 0.49816 | 0 | -0.08869 | 0.01829 | 0 | 13.11096 | Site_A | 12.64941 |
| 23 | 17 | 813.62387 | 0.1393 | 0.07785 | 0.07358 | 2.44151 | 0.51893 | 0 | -0.09178 | 0.0188 | 0 | 13.30114 | Site_A | 13.91341 |
| 26 | 17 | 816.96759 | 0.13835 | 0.07851 | 0.07805 | 2.44328 | 0.51785 | 0 | -0.09074 | 0.01854 | 0 | 13.46383 | Site_A | 17.25713 |
| 29 | 17 | 820.39579 | 0.13981 | 0.07942 | 0.07837 | 2.44739 | 0.51833 | 0 | -0.08982 | 0.01835 | 0 | 13.62416 | Site_A | 20.68532 |
| 32 | 17 | 822.11258 | 0.14019 | 0.08055 | 0.0818 | 2.52412 | 0.52962 | 0 | -0.09174 | 0.01858 | 0 | 13.75716 | Site_A | 22.40212 |
| 35 | 17 | 824.41081 | 0.13821 | 0.08094 | 0.08772 | 2.55731 | 0.53689 | 0 | -0.09225 | 0.0187 | 0 | 13.86093 | Site_A | 24.70035 |
| 2 | 20 | 846.9174 | 0.29349 | 0.08379 | 0.00046 | 0.43676 | 0.15691 | 0.00538 | -0.01874 | 0.00623 | 0.00262 | 11.65117 | Site_A | 47.20694 |
| 5 | 20 | 829.58716 | 0.27373 | 0.08169 | 0.00081 | 1.04367 | 0.26246 | 0.00007 | -0.04198 | 0.01013 | 0.00003 | 12.42951 | Site_A | 29.8767 |
| 8 | 20 | 824.10913 | 0.26683 | 0.08149 | 0.00106 | 1.26227 | 0.30409 | 0.00003 | -0.05084 | 0.01171 | 0.00001 | 12.41333 | Site_A | 24.39866 |
| 11 | 20 | 814.99601 | 0.26073 | 0.08189 | 0.00145 | 1.69018 | 0.38614 | 0.00001 | -0.06685 | 0.01467 | 0.00001 | 12.64095 | Site_A | 15.28555 |
| 14 | 20 | 809.71755 | 0.25157 | 0.08256 | 0.00231 | 2.03569 | 0.44815 | 0.00001 | -0.07935 | 0.01684 | 0 | 12.82659 | Site_A | 10.00709 |
| 17 | 20 | 808.38522 | 0.24311 | 0.08334 | 0.00353 | 2.19222 | 0.4785 | 0 | -0.08465 | 0.01779 | 0 | 12.94825 | Site_A | 8.67476 |
| 20 | 20 | 808.24357 | 0.23263 | 0.08334 | 0.00525 | 2.3436 | 0.51065 | 0 | -0.08925 | 0.01872 | 0 | 13.1299 | Site_A | 8.53311 |
| 23 | 20 | 809.68139 | 0.22686 | 0.0835 | 0.00659 | 2.44062 | 0.52822 | 0 | -0.09167 | 0.01911 | 0 | 13.31138 | Site_A | 9.97093 |
| 26 | 20 | 813.10783 | 0.22532 | 0.08402 | 0.00733 | 2.42653 | 0.52438 | 0 | -0.09008 | 0.01874 | 0 | 13.46882 | Site_A | 13.39737 |
| 29 | 20 | 816.58705 | 0.22665 | 0.08486 | 0.00757 | 2.41871 | 0.52269 | 0 | -0.08874 | 0.01847 | 0 | 13.62737 | Site_A | 16.87659 |
| 32 | 20 | 818.31331 | 0.22867 | 0.08607 | 0.00789 | 2.48623 | 0.5326 | 0 | -0.09034 | 0.01865 | 0 | 13.75973 | Site_A | 18.60285 |
| 35 | 20 | 820.67802 | 0.22707 | 0.08659 | 0.00873 | 2.50796 | 0.53856 | 0 | -0.09047 | 0.01872 | 0 | 13.86011 | Site_A | 20.96756 |
| 2 | 23 | 844.78951 | 0.35512 | 0.09457 | 0.00017 | 0.43394 | 0.15676 | 0.00564 | -0.01878 | 0.00621 | 0.00248 | 11.55272 | Site_A | 45.07905 |
| 5 | 23 | 827.77383 | 0.32981 | 0.0922 | 0.00035 | 1.03499 | 0.26179 | 0.00008 | -0.04179 | 0.0101 | 0.00003 | 12.38249 | Site_A | 28.06337 |
| 8 | 23 | 822.41377 | 0.31938 | 0.09139 | 0.00047 | 1.25108 | 0.30386 | 0.00004 | -0.05056 | 0.0117 | 0.00002 | 12.37144 | Site_A | 22.70331 |
| 11 | 23 | 813.36099 | 0.31061 | 0.0911 | 0.00065 | 1.68757 | 0.38781 | 0.00001 | -0.06686 | 0.01473 | 0.00001 | 12.61986 | Site_A | 13.65053 |
| 14 | 23 | 807.95797 | 0.30281 | 0.09159 | 0.00095 | 2.04979 | 0.45323 | 0.00001 | -0.07994 | 0.01703 | 0 | 12.82129 | Site_A | 8.24751 |
| 17 | 23 | 806.37579 | 0.29918 | 0.09258 | 0.00123 | 2.23512 | 0.48928 | 0 | -0.08624 | 0.01817 | 0 | 12.95856 | Site_A | 6.66533 |
| 20 | 23 | 806.26465 | 0.28827 | 0.09241 | 0.00181 | 2.40222 | 0.52377 | 0 | -0.09136 | 0.01918 | 0 | 13.14683 | Site_A | 6.55419 |
| 23 | 23 | 807.96091 | 0.27787 | 0.09234 | 0.00262 | 2.48817 | 0.53912 | 0 | -0.09336 | 0.01949 | 0 | 13.32618 | Site_A | 8.25045 |
| 26 | 23 | 811.54688 | 0.2728 | 0.09246 | 0.00317 | 2.4579 | 0.5323 | 0 | -0.09118 | 0.01901 | 0 | 13.47884 | Site_A | 11.83642 |
| 29 | 23 | 815.10207 | 0.27257 | 0.09313 | 0.00342 | 2.44347 | 0.52975 | 0 | -0.0896 | 0.0187 | 0 | 13.63539 | Site_A | 15.39161 |
| 32 | 23 | 816.90499 | 0.27369 | 0.09434 | 0.00372 | 2.50899 | 0.53954 | 0 | -0.09112 | 0.01888 | 0 | 13.76742 | Site_A | 17.19453 |
| 35 | 23 | 819.30592 | 0.27261 | 0.09521 | 0.00419 | 2.52578 | 0.54501 | 0 | -0.09106 | 0.01893 | 0 | 13.86805 | Site_A | 19.59546 |
| 2 | 26 | 839.82954 | 0.45503 | 0.10644 | 0.00002 | 0.42385 | 0.15711 | 0.00698 | -0.01855 | 0.0062 | 0.00276 | 11.42601 | Site_A | 40.11908 |
| 5 | 26 | 822.97386 | 0.4284 | 0.10393 | 0.00004 | 1.02078 | 0.26118 | 0.00009 | -0.04133 | 0.01003 | 0.00004 | 12.34995 | Site_A | 23.2634 |
| 8 | 26 | 817.74353 | 0.41559 | 0.10304 | 0.00005 | 1.2351 | 0.30422 | 0.00005 | -0.05009 | 0.01168 | 0.00002 | 12.32982 | Site_A | 18.03307 |
| 11 | 26 | 808.75686 | 0.40442 | 0.10208 | 0.00007 | 1.68183 | 0.3914 | 0.00002 | -0.0668 | 0.01485 | 0.00001 | 12.58801 | Site_A | 9.0464 |
| 14 | 26 | 803.29386 | 0.3975 | 0.10222 | 0.0001 | 2.06517 | 0.46195 | 0.00001 | -0.08064 | 0.01735 | 0 | 12.80442 | Site_A | 3.5834 |
| 17 | 26 | 801.63096 | 0.39655 | 0.10336 | 0.00012 | 2.26961 | 0.50274 | 0.00001 | -0.08759 | 0.01866 | 0 | 12.9554 | Site_A | 1.9205 |
| 20 | 26 | 801.33159 | 0.38952 | 0.10317 | 0.00016 | 2.46995 | 0.54321 | 0.00001 | -0.09385 | 0.01988 | 0 | 13.15839 | Site_A | 1.62113 |
| 23 | 26 | 803.08475 | 0.37865 | 0.10267 | 0.00023 | 2.56749 | 0.55934 | 0 | -0.09621 | 0.02021 | 0 | 13.34257 | Site_A | 3.37429 |
| 26 | 26 | 807.06476 | 0.36745 | 0.10235 | 0.00033 | 2.50814 | 0.54737 | 0 | -0.09296 | 0.01954 | 0 | 13.49041 | Site_A | 7.3543 |
| 29 | 26 | 810.81403 | 0.3637 | 0.10253 | 0.00039 | 2.47041 | 0.54106 | 0 | -0.09055 | 0.01908 | 0 | 13.64118 | Site_A | 11.10357 |
| 32 | 26 | 812.70151 | 0.36417 | 0.10355 | 0.00044 | 2.52891 | 0.55017 | 0 | -0.09182 | 0.01922 | 0 | 13.77151 | Site_A | 12.99105 |
| 35 | 26 | 815.21934 | 0.36254 | 0.10448 | 0.00052 | 2.53962 | 0.5554 | 0 | -0.09154 | 0.01926 | 0 | 13.87185 | Site_A | 15.50888 |
| 2 | 29 | 840.16357 | 0.49816 | 0.11971 | 0.00003 | 0.41651 | 0.15583 | 0.00752 | -0.01839 | 0.00617 | 0.00287 | 11.32696 | Site_A | 40.45311 |
| 5 | 29 | 823.49778 | 0.46872 | 0.11734 | 0.00006 | 1.00743 | 0.25895 | 0.0001 | -0.0409 | 0.00995 | 0.00004 | 12.31665 | Site_A | 23.78732 |
| 8 | 29 | 817.89568 | 0.46139 | 0.11665 | 0.00008 | 1.22548 | 0.30219 | 0.00005 | -0.04975 | 0.01159 | 0.00002 | 12.31595 | Site_A | 18.18522 |
| 11 | 29 | 808.7458 | 0.4506 | 0.11537 | 0.00009 | 1.67901 | 0.3904 | 0.00002 | -0.06674 | 0.0148 | 0.00001 | 12.57891 | Site_A | 9.03534 |
| 14 | 29 | 803.14681 | 0.4432 | 0.11494 | 0.00012 | 2.06967 | 0.46295 | 0.00001 | -0.08091 | 0.01738 | 0 | 12.79 | Site_A | 3.43635 |
| 17 | 29 | 801.4572 | 0.44172 | 0.11587 | 0.00014 | 2.28157 | 0.50569 | 0.00001 | -0.08812 | 0.01877 | 0 | 12.94509 | Site_A | 1.74674 |
| 20 | 29 | 801.03059 | 0.43632 | 0.11582 | 0.00016 | 2.50103 | 0.55044 | 0.00001 | -0.09503 | 0.02013 | 0 | 13.15983 | Site_A | 1.32013 |
| 23 | 29 | 802.43381 | 0.42969 | 0.11522 | 0.00019 | 2.63486 | 0.57195 | 0 | -0.09865 | 0.02066 | 0 | 13.35392 | Site_A | 2.72335 |
| 26 | 29 | 806.38201 | 0.41694 | 0.11438 | 0.00027 | 2.58781 | 0.56027 | 0 | -0.0958 | 0.02 | 0 | 13.50577 | Site_A | 6.67155 |
| 29 | 29 | 810.38531 | 0.40739 | 0.11406 | 0.00035 | 2.53724 | 0.55172 | 0 | -0.09289 | 0.01946 | 0 | 13.6565 | Site_A | 10.67484 |
| 32 | 29 | 812.37614 | 0.40458 | 0.11465 | 0.00042 | 2.58612 | 0.55905 | 0 | -0.0938 | 0.01953 | 0 | 13.78539 | Site_A | 12.66568 |
| 35 | 29 | 814.9726 | 0.40102 | 0.11555 | 0.00052 | 2.59373 | 0.56383 | 0 | -0.09339 | 0.01954 | 0 | 13.88687 | Site_A | 15.26214 |
| 2 | 32 | 839.03748 | 0.56034 | 0.13337 | 0.00003 | 0.42265 | 0.15556 | 0.00659 | -0.01869 | 0.00619 | 0.00253 | 11.30567 | Site_A | 39.32702 |
| 5 | 32 | 822.77943 | 0.52422 | 0.13122 | 0.00006 | 0.99558 | 0.25626 | 0.0001 | -0.0406 | 0.00988 | 0.00004 | 12.26199 | Site_A | 23.06897 |
| 8 | 32 | 817.15777 | 0.52075 | 0.13158 | 0.00008 | 1.21703 | 0.30085 | 0.00005 | -0.04955 | 0.01156 | 0.00002 | 12.27994 | Site_A | 17.44731 |
| 11 | 32 | 807.71726 | 0.51587 | 0.13078 | 0.00008 | 1.68208 | 0.39157 | 0.00002 | -0.06695 | 0.01486 | 0.00001 | 12.56256 | Site_A | 8.0068 |
| 14 | 32 | 801.86077 | 0.51212 | 0.13036 | 0.00009 | 2.09465 | 0.46887 | 0.00001 | -0.08193 | 0.01761 | 0 | 12.78291 | Site_A | 2.15031 |
| 17 | 32 | 800.1135 | 0.50973 | 0.13071 | 0.0001 | 2.31604 | 0.51404 | 0.00001 | -0.08952 | 0.01909 | 0 | 12.93592 | Site_A | 0.40303 |
| 20 | 32 | 799.71046 | 0.50322 | 0.13049 | 0.00012 | 2.54588 | 0.56124 | 0.00001 | -0.09677 | 0.02054 | 0 | 13.15458 | Site_A | 0 |
| 23 | 32 | 800.98971 | 0.49836 | 0.12994 | 0.00013 | 2.69871 | 0.58619 | 0 | -0.10103 | 0.02118 | 0 | 13.35592 | Site_A | 1.27925 |
| 26 | 32 | 804.70995 | 0.48776 | 0.12873 | 0.00015 | 2.66903 | 0.57577 | 0 | -0.09877 | 0.02057 | 0 | 13.51106 | Site_A | 4.99949 |
| 29 | 32 | 808.73743 | 0.47608 | 0.12784 | 0.0002 | 2.6203 | 0.5659 | 0 | -0.09588 | 0.01997 | 0 | 13.6642 | Site_A | 9.02697 |
| 32 | 32 | 810.93052 | 0.46826 | 0.1279 | 0.00025 | 2.65979 | 0.57169 | 0 | -0.0964 | 0.01998 | 0 | 13.79554 | Site_A | 11.22006 |
| 35 | 32 | 813.64158 | 0.46104 | 0.12826 | 0.00032 | 2.6557 | 0.57461 | 0 | -0.09554 | 0.01992 | 0 | 13.89772 | Site_A | 13.93112 |
| 2 | 35 | 842.28199 | 0.54667 | 0.14259 | 0.00013 | 0.44094 | 0.15628 | 0.00478 | -0.0193 | 0.00622 | 0.00192 | 11.42369 | Site_A | 42.57153 |
| 5 | 35 | 825.23455 | 0.52489 | 0.14232 | 0.00023 | 1.02281 | 0.25601 | 0.00006 | -0.04166 | 0.00991 | 0.00003 | 12.27663 | Site_A | 25.52409 |
| 8 | 35 | 819.76059 | 0.51901 | 0.14304 | 0.00029 | 1.23977 | 0.29886 | 0.00003 | -0.05049 | 0.01154 | 0.00001 | 12.2785 | Site_A | 20.05013 |
| 11 | 35 | 810.3523 | 0.51757 | 0.14382 | 0.00032 | 1.70078 | 0.38835 | 0.00001 | -0.06773 | 0.01479 | 0 | 12.55515 | Site_A | 10.64184 |
| 14 | 35 | 804.25175 | 0.52107 | 0.1446 | 0.00031 | 2.11652 | 0.46676 | 0.00001 | -0.0828 | 0.01757 | 0 | 12.78104 | Site_A | 4.54129 |
| 17 | 35 | 802.34839 | 0.52053 | 0.14497 | 0.00033 | 2.34926 | 0.51393 | 0 | -0.09078 | 0.01911 | 0 | 12.9391 | Site_A | 2.63793 |
| 20 | 35 | 801.94298 | 0.512 | 0.14435 | 0.00039 | 2.5798 | 0.56125 | 0 | -0.09807 | 0.02056 | 0 | 13.15247 | Site_A | 2.23252 |
| 23 | 35 | 803.32438 | 0.50389 | 0.14352 | 0.00045 | 2.73239 | 0.58641 | 0 | -0.10231 | 0.02121 | 0 | 13.35307 | Site_A | 3.61392 |
| 26 | 35 | 806.93166 | 0.49352 | 0.14199 | 0.00051 | 2.71889 | 0.57885 | 0 | -0.1006 | 0.02069 | 0 | 13.51349 | Site_A | 7.2212 |
| 29 | 35 | 810.75138 | 0.48308 | 0.1406 | 0.00059 | 2.68981 | 0.57092 | 0 | -0.09841 | 0.02018 | 0 | 13.66681 | Site_A | 11.04092 |
| 32 | 35 | 812.91563 | 0.47276 | 0.14002 | 0.00073 | 2.74034 | 0.577 | 0 | -0.0993 | 0.0202 | 0 | 13.79828 | Site_A | 13.20516 |
| 35 | 35 | 815.7559 | 0.45908 | 0.13944 | 0.00099 | 2.73623 | 0.57939 | 0 | -0.09839 | 0.02011 | 0 | 13.9054 | Site_A | 16.04544 |
| 2 | 2 | 805.6533 | 0.0317 | 0.02685 | 0.23777 | 0.72083 | 0.21821 | 0.00096 | -0.02779 | 0.00828 | 0.00079 | 12.96905 | Site_B | 53.81078 |
| 5 | 2 | 783.7122 | 0.03931 | 0.02702 | 0.14576 | 1.46788 | 0.30456 | 0 | -0.0553 | 0.01122 | 0 | 13.27185 | Site_B | 31.86969 |
| 8 | 2 | 778.1719 | 0.04057 | 0.02707 | 0.13395 | 1.73671 | 0.34699 | 0 | -0.06591 | 0.01278 | 0 | 13.17462 | Site_B | 26.32938 |
| 11 | 2 | 768.36085 | 0.03784 | 0.02757 | 0.16993 | 2.16674 | 0.4116 | 0 | -0.08212 | 0.01508 | 0 | 13.19172 | Site_B | 16.51833 |
| 14 | 2 | 764.13923 | 0.03657 | 0.02779 | 0.18815 | 2.38653 | 0.44053 | 0 | -0.09044 | 0.01608 | 0 | 13.19374 | Site_B | 12.29671 |
| 17 | 2 | 763.79213 | 0.03659 | 0.0277 | 0.18651 | 2.47633 | 0.45649 | 0 | -0.09356 | 0.01653 | 0 | 13.23398 | Site_B | 11.94961 |
| 20 | 2 | 762.96765 | 0.03505 | 0.02786 | 0.20832 | 2.67882 | 0.49457 | 0 | -0.10001 | 0.01767 | 0 | 13.39216 | Site_B | 11.12513 |
| 23 | 2 | 763.28818 | 0.03531 | 0.02797 | 0.20682 | 2.85995 | 0.52466 | 0 | -0.10544 | 0.01853 | 0 | 13.56167 | Site_B | 11.44566 |
| 26 | 2 | 765.40996 | 0.03673 | 0.02816 | 0.19215 | 2.96948 | 0.54098 | 0 | -0.10794 | 0.01888 | 0 | 13.7546 | Site_B | 13.56744 |
| 29 | 2 | 769.28783 | 0.03842 | 0.02834 | 0.17525 | 3.00595 | 0.54888 | 0 | -0.10775 | 0.01894 | 0 | 13.94854 | Site_B | 17.44531 |
| 32 | 2 | 770.74121 | 0.03903 | 0.02841 | 0.16954 | 3.111 | 0.56381 | 0 | -0.11047 | 0.01929 | 0 | 14.08067 | Site_B | 18.89869 |
| 35 | 2 | 773.36372 | 0.03959 | 0.02823 | 0.16079 | 3.14746 | 0.57244 | 0 | -0.11089 | 0.01943 | 0 | 14.1921 | Site_B | 21.5212 |
| 2 | 5 | 802.48969 | 0.08022 | 0.03572 | 0.02472 | 0.6991 | 0.21969 | 0.00146 | -0.02685 | 0.00835 | 0.00131 | 13.01943 | Site_B | 50.64717 |
| 5 | 5 | 781.03711 | 0.08182 | 0.03541 | 0.02086 | 1.45171 | 0.30731 | 0 | -0.05462 | 0.01132 | 0 | 13.28815 | Site_B | 29.19459 |
| 8 | 5 | 774.95014 | 0.08885 | 0.03581 | 0.0131 | 1.71241 | 0.34747 | 0 | -0.06501 | 0.01279 | 0 | 13.17131 | Site_B | 23.10762 |
| 11 | 5 | 765.14113 | 0.08731 | 0.03655 | 0.01689 | 2.13487 | 0.41122 | 0 | -0.08098 | 0.01506 | 0 | 13.18163 | Site_B | 13.29861 |
| 14 | 5 | 761.22623 | 0.08356 | 0.03676 | 0.02304 | 2.34152 | 0.43831 | 0 | -0.08881 | 0.01598 | 0 | 13.18216 | Site_B | 9.38371 |
| 17 | 5 | 760.71283 | 0.08591 | 0.03699 | 0.02021 | 2.44308 | 0.45643 | 0 | -0.09235 | 0.01652 | 0 | 13.22695 | Site_B | 8.87031 |
| 20 | 5 | 759.76027 | 0.08637 | 0.03728 | 0.02052 | 2.6587 | 0.49684 | 0 | -0.0993 | 0.01775 | 0 | 13.3876 | Site_B | 7.91775 |
| 23 | 5 | 760.12249 | 0.08732 | 0.03797 | 0.02147 | 2.83822 | 0.52618 | 0 | -0.1047 | 0.01859 | 0 | 13.55447 | Site_B | 8.27997 |
| 26 | 5 | 762.10577 | 0.09057 | 0.03839 | 0.01832 | 2.9398 | 0.53984 | 0 | -0.10694 | 0.01885 | 0 | 13.74507 | Site_B | 10.26325 |
| 29 | 5 | 765.79025 | 0.09481 | 0.03896 | 0.01496 | 2.96567 | 0.54508 | 0 | -0.10638 | 0.0188 | 0 | 13.93948 | Site_B | 13.94774 |
| 32 | 5 | 767.11868 | 0.09685 | 0.03917 | 0.01342 | 3.06568 | 0.55842 | 0 | -0.10895 | 0.01911 | 0 | 14.06926 | Site_B | 15.27617 |
| 35 | 5 | 769.61875 | 0.0981 | 0.0389 | 0.01167 | 3.09915 | 0.56625 | 0 | -0.10932 | 0.01923 | 0 | 14.1753 | Site_B | 17.77624 |
| 2 | 8 | 803.37498 | 0.09038 | 0.04495 | 0.04438 | 0.71644 | 0.22941 | 0.00179 | -0.02748 | 0.00872 | 0.00162 | 13.03409 | Site_B | 51.53246 |
| 5 | 8 | 782.33207 | 0.08604 | 0.04448 | 0.05309 | 1.47546 | 0.30955 | 0 | -0.05542 | 0.0114 | 0 | 13.31219 | Site_B | 30.48955 |
| 8 | 8 | 777.05534 | 0.08485 | 0.04519 | 0.06041 | 1.73609 | 0.35118 | 0 | -0.06574 | 0.01293 | 0 | 13.20485 | Site_B | 25.21282 |
| 11 | 8 | 767.15248 | 0.08328 | 0.04628 | 0.07195 | 2.15653 | 0.41411 | 0 | -0.08163 | 0.01516 | 0 | 13.20868 | Site_B | 15.30996 |
| 14 | 8 | 763.19116 | 0.07725 | 0.04642 | 0.09612 | 2.35876 | 0.44032 | 0 | -0.08934 | 0.01605 | 0 | 13.20049 | Site_B | 11.34864 |
| 17 | 8 | 762.89349 | 0.07699 | 0.04666 | 0.09892 | 2.4454 | 0.45586 | 0 | -0.09235 | 0.01649 | 0 | 13.2399 | Site_B | 11.05097 |
| 20 | 8 | 761.88806 | 0.07829 | 0.04697 | 0.09553 | 2.65867 | 0.49576 | 0 | -0.0992 | 0.0177 | 0 | 13.40053 | Site_B | 10.04554 |
| 23 | 8 | 762.26746 | 0.07829 | 0.04754 | 0.09961 | 2.84023 | 0.5262 | 0 | -0.10466 | 0.01858 | 0 | 13.56822 | Site_B | 10.42494 |
| 26 | 8 | 764.46201 | 0.07966 | 0.04824 | 0.09866 | 2.94292 | 0.54167 | 0 | -0.10695 | 0.01891 | 0 | 13.75777 | Site_B | 12.61949 |
| 29 | 8 | 768.2835 | 0.08363 | 0.04894 | 0.08748 | 2.9662 | 0.54752 | 0 | -0.10631 | 0.01889 | 0 | 13.95087 | Site_B | 16.44098 |
| 32 | 8 | 769.60208 | 0.08754 | 0.04958 | 0.07747 | 3.06372 | 0.56108 | 0 | -0.10877 | 0.01919 | 0 | 14.08358 | Site_B | 17.75956 |
| 35 | 8 | 772.12636 | 0.08958 | 0.04918 | 0.06854 | 3.09437 | 0.56853 | 0 | -0.10901 | 0.0193 | 0 | 14.1937 | Site_B | 20.28384 |
| 2 | 11 | 800.45431 | 0.13669 | 0.05132 | 0.00773 | 0.72071 | 0.22851 | 0.00161 | -0.02765 | 0.00869 | 0.00146 | 13.03235 | Site_B | 48.61179 |
| 5 | 11 | 779.08252 | 0.1356 | 0.0485 | 0.00517 | 1.49599 | 0.31268 | 0 | -0.05614 | 0.01151 | 0 | 13.32342 | Site_B | 27.24 |
| 8 | 11 | 774.1745 | 0.13085 | 0.04926 | 0.00789 | 1.7626 | 0.35638 | 0 | -0.06669 | 0.01311 | 0 | 13.21582 | Site_B | 22.33198 |
| 11 | 11 | 764.8251 | 0.12367 | 0.05051 | 0.01434 | 2.17722 | 0.41951 | 0 | -0.08234 | 0.01533 | 0 | 13.22079 | Site_B | 12.98258 |
| 14 | 11 | 761.07039 | 0.11636 | 0.05088 | 0.02221 | 2.36234 | 0.44285 | 0 | -0.08942 | 0.01612 | 0 | 13.20922 | Site_B | 9.22787 |
| 17 | 11 | 760.87754 | 0.1146 | 0.05101 | 0.02467 | 2.44294 | 0.45734 | 0 | -0.09221 | 0.01653 | 0 | 13.24617 | Site_B | 9.03502 |
| 20 | 11 | 759.96222 | 0.11433 | 0.05115 | 0.02539 | 2.65462 | 0.49672 | 0 | -0.09901 | 0.01772 | 0 | 13.40532 | Site_B | 8.1197 |
| 23 | 11 | 760.31175 | 0.1151 | 0.05166 | 0.02589 | 2.83389 | 0.52617 | 0 | -0.10443 | 0.01858 | 0 | 13.56867 | Site_B | 8.46923 |
| 26 | 11 | 762.52746 | 0.11619 | 0.05223 | 0.02612 | 2.93384 | 0.54142 | 0 | -0.10663 | 0.01889 | 0 | 13.7573 | Site_B | 10.68494 |
| 29 | 11 | 766.44775 | 0.11879 | 0.053 | 0.025 | 2.95179 | 0.547 | 0 | -0.1058 | 0.01886 | 0 | 13.94999 | Site_B | 14.60523 |
| 32 | 11 | 767.78067 | 0.12263 | 0.05379 | 0.02261 | 3.04374 | 0.55978 | 0 | -0.10808 | 0.01914 | 0 | 14.08151 | Site_B | 15.93815 |
| 35 | 11 | 770.25546 | 0.12544 | 0.05369 | 0.01948 | 3.06845 | 0.56638 | 0 | -0.10811 | 0.01922 | 0 | 14.19127 | Site_B | 18.41295 |
| 2 | 14 | 801.24657 | 0.14253 | 0.05783 | 0.01373 | 0.72953 | 0.23607 | 0.002 | -0.02807 | 0.00897 | 0.00175 | 12.99283 | Site_B | 49.40405 |
| 5 | 14 | 779.53552 | 0.14553 | 0.05474 | 0.00784 | 1.50902 | 0.31301 | 0 | -0.05669 | 0.01151 | 0 | 13.30915 | Site_B | 27.693 |
| 8 | 14 | 774.35682 | 0.14332 | 0.05523 | 0.00946 | 1.78527 | 0.3586 | 0 | -0.06755 | 0.01318 | 0 | 13.21448 | Site_B | 22.5143 |
| 11 | 14 | 765.02921 | 0.13461 | 0.05639 | 0.01698 | 2.21208 | 0.42424 | 0 | -0.08363 | 0.0155 | 0 | 13.22564 | Site_B | 13.18669 |
| 14 | 14 | 761.53512 | 0.12251 | 0.05689 | 0.03127 | 2.39345 | 0.4471 | 0 | -0.09054 | 0.01627 | 0 | 13.21694 | Site_B | 9.6926 |
| 17 | 14 | 761.38516 | 0.12005 | 0.0572 | 0.03583 | 2.46605 | 0.45996 | 0 | -0.09304 | 0.01661 | 0 | 13.2527 | Site_B | 9.54264 |
| 20 | 14 | 760.52428 | 0.11846 | 0.05719 | 0.03832 | 2.67485 | 0.49864 | 0 | -0.09972 | 0.01778 | 0 | 13.41132 | Site_B | 8.68177 |
| 23 | 14 | 760.99098 | 0.11674 | 0.05747 | 0.04223 | 2.85244 | 0.52765 | 0 | -0.10506 | 0.01861 | 0 | 13.57509 | Site_B | 9.14846 |
| 26 | 14 | 763.27267 | 0.11643 | 0.05789 | 0.0443 | 2.95223 | 0.54314 | 0 | -0.10726 | 0.01894 | 0 | 13.76218 | Site_B | 11.43015 |
| 29 | 14 | 767.29579 | 0.11712 | 0.05851 | 0.04533 | 2.97376 | 0.55003 | 0 | -0.10655 | 0.01896 | 0 | 13.95478 | Site_B | 15.45327 |
| 32 | 14 | 768.82346 | 0.11828 | 0.05945 | 0.04665 | 3.06579 | 0.56373 | 0 | -0.10882 | 0.01926 | 0 | 14.08692 | Site_B | 16.98094 |
| 35 | 14 | 771.40679 | 0.12047 | 0.05969 | 0.04355 | 3.08881 | 0.57038 | 0 | -0.10878 | 0.01934 | 0 | 14.19803 | Site_B | 19.56427 |
| 2 | 17 | 799.95028 | 0.17078 | 0.06381 | 0.00744 | 0.73404 | 0.22677 | 0.00121 | -0.02833 | 0.00861 | 0.001 | 12.95612 | Site_B | 48.10776 |
| 5 | 17 | 778.09602 | 0.1743 | 0.05856 | 0.00291 | 1.52093 | 0.31371 | 0 | -0.0572 | 0.01153 | 0 | 13.294 | Site_B | 26.2535 |
| 8 | 17 | 772.82955 | 0.17275 | 0.05882 | 0.00332 | 1.80199 | 0.36085 | 0 | -0.06821 | 0.01325 | 0 | 13.20853 | Site_B | 20.98703 |
| 11 | 17 | 763.23713 | 0.16897 | 0.06024 | 0.00503 | 2.24796 | 0.43047 | 0 | -0.08496 | 0.01571 | 0 | 13.23017 | Site_B | 11.39462 |
| 14 | 17 | 759.9291 | 0.15553 | 0.0608 | 0.01053 | 2.43059 | 0.45358 | 0 | -0.0919 | 0.01648 | 0 | 13.22478 | Site_B | 8.08659 |
| 17 | 17 | 760.12474 | 0.14806 | 0.06105 | 0.0153 | 2.49354 | 0.46539 | 0 | -0.09403 | 0.01679 | 0 | 13.25895 | Site_B | 8.28222 |
| 20 | 17 | 759.35228 | 0.14504 | 0.06096 | 0.01734 | 2.69522 | 0.50267 | 0 | -0.10046 | 0.01791 | 0 | 13.41503 | Site_B | 7.50976 |
| 23 | 17 | 759.90468 | 0.14171 | 0.06104 | 0.02025 | 2.86905 | 0.53077 | 0 | -0.10565 | 0.01871 | 0 | 13.57745 | Site_B | 8.06216 |
| 26 | 17 | 762.31036 | 0.13914 | 0.06124 | 0.02309 | 2.96593 | 0.54579 | 0 | -0.10774 | 0.01902 | 0 | 13.76421 | Site_B | 10.46784 |
| 29 | 17 | 766.40616 | 0.13878 | 0.06183 | 0.02479 | 2.98385 | 0.55262 | 0 | -0.1069 | 0.01904 | 0 | 13.95594 | Site_B | 14.56364 |
| 32 | 17 | 767.97726 | 0.13946 | 0.06279 | 0.02634 | 3.07635 | 0.56687 | 0 | -0.10918 | 0.01936 | 0 | 14.08869 | Site_B | 16.13474 |
| 35 | 17 | 770.68291 | 0.13989 | 0.06314 | 0.02673 | 3.0972 | 0.57376 | 0 | -0.10906 | 0.01944 | 0 | 14.20014 | Site_B | 18.84039 |
| 2 | 20 | 796.13476 | 0.23227 | 0.07364 | 0.00161 | 0.72557 | 0.20481 | 0.0004 | -0.02818 | 0.00778 | 0.00029 | 12.87408 | Site_B | 44.29224 |
| 5 | 20 | 774.19159 | 0.22708 | 0.06114 | 0.0002 | 1.52773 | 0.31482 | 0 | -0.05763 | 0.01157 | 0 | 13.25559 | Site_B | 22.34907 |
| 8 | 20 | 768.84448 | 0.22584 | 0.06117 | 0.00022 | 1.81523 | 0.36305 | 0 | -0.06884 | 0.01334 | 0 | 13.18357 | Site_B | 17.00197 |
| 11 | 20 | 759.10789 | 0.2249 | 0.06244 | 0.00032 | 2.27719 | 0.43589 | 0 | -0.08614 | 0.0159 | 0 | 13.21855 | Site_B | 7.26537 |
| 14 | 20 | 755.70728 | 0.216 | 0.06325 | 0.00064 | 2.47486 | 0.46222 | 0 | -0.09357 | 0.01677 | 0 | 13.22487 | Site_B | 3.86476 |
| 17 | 20 | 756.0711 | 0.2082 | 0.06353 | 0.00105 | 2.54165 | 0.47523 | 0 | -0.09583 | 0.01712 | 0 | 13.26179 | Site_B | 4.22858 |
| 20 | 20 | 755.69553 | 0.20017 | 0.06323 | 0.00155 | 2.73421 | 0.51175 | 0 | -0.1019 | 0.01822 | 0 | 13.4155 | Site_B | 3.85302 |
| 23 | 20 | 756.42007 | 0.1958 | 0.07184 | 0.00642 | 2.89667 | 0.55305 | 0 | -0.10668 | 0.01969 | 0 | 13.57585 | Site_B | 4.57755 |
| 26 | 20 | 758.96619 | 0.19369 | 0.07244 | 0.0075 | 2.98769 | 0.56558 | 0 | -0.10854 | 0.01992 | 0 | 13.76339 | Site_B | 7.12367 |
| 29 | 20 | 763.18747 | 0.19364 | 0.07375 | 0.00865 | 2.99975 | 0.5703 | 0 | -0.10746 | 0.01986 | 0 | 13.95748 | Site_B | 11.34496 |
| 32 | 20 | 764.79152 | 0.19711 | 0.07521 | 0.00877 | 3.09088 | 0.58413 | 0 | -0.10966 | 0.02017 | 0 | 14.09364 | Site_B | 12.94901 |
| 35 | 20 | 767.50895 | 0.19936 | 0.07547 | 0.00825 | 3.11222 | 0.59073 | 0 | -0.10953 | 0.02025 | 0 | 14.20706 | Site_B | 15.66643 |
| 2 | 23 | 792.78361 | 0.29737 | 0.08614 | 0.00056 | 0.7222 | 0.19576 | 0.00022 | -0.02823 | 0.00743 | 0.00014 | 12.79214 | Site_B | 40.94109 |
| 5 | 23 | 771.2624 | 0.28464 | 0.09025 | 0.00161 | 1.5143 | 0.42217 | 0.00033 | -0.05726 | 0.01571 | 0.00027 | 13.22313 | Site_B | 19.41989 |
| 8 | 23 | 765.9313 | 0.27623 | 0.18015 | 0.1252 | 1.81077 | 1.35848 | 0.18255 | -0.06883 | 0.05104 | 0.17745 | 13.15346 | Site_B | 14.08878 |
| 11 | 23 | 756.07804 | 0.2785 | 0.10346 | 0.00711 | 2.29072 | 0.79251 | 0.00385 | -0.08677 | 0.02945 | 0.00322 | 13.20034 | Site_B | 4.23552 |
| 14 | 23 | 752.45471 | 0.27496 | 0.09258 | 0.00298 | 2.51069 | 0.65815 | 0.00014 | -0.09498 | 0.02421 | 0.00009 | 13.21643 | Site_B | 0.61219 |
| 17 | 23 | 752.51956 | 0.27403 | 0.09055 | 0.00248 | 2.60444 | 0.62835 | 0.00003 | -0.09818 | 0.0229 | 0.00002 | 13.26364 | Site_B | 0.67704 |
| 20 | 23 | 752.2357 | 0.26635 | 0.09069 | 0.00331 | 2.81485 | 0.69229 | 0.00005 | -0.10487 | 0.02495 | 0.00003 | 13.42093 | Site_B | 0.39318 |
| 23 | 23 | 753.29208 | 0.25875 | 0.0916 | 0.00473 | 2.97038 | 0.73377 | 0.00005 | -0.10935 | 0.0262 | 0.00003 | 13.58143 | Site_B | 1.44957 |
| 26 | 23 | 755.98701 | 0.25363 | 0.09516 | 0.00769 | 3.04248 | 0.79881 | 0.00014 | -0.1105 | 0.02825 | 0.00009 | 13.76709 | Site_B | 4.1445 |
| 29 | 23 | 760.31867 | 0.25231 | 0.10002 | 0.01165 | 3.04425 | 0.857 | 0.00038 | -0.10902 | 0.03001 | 0.00028 | 13.96173 | Site_B | 8.47615 |
| 32 | 23 | 761.99131 | 0.25538 | 0.09919 | 0.01003 | 3.13477 | 0.83971 | 0.00019 | -0.11117 | 0.02914 | 0.00014 | 14.09878 | Site_B | 10.14879 |
| 35 | 23 | 764.74875 | 0.25797 | 0.09812 | 0.00856 | 3.15326 | 0.8229 | 0.00013 | -0.11093 | 0.02831 | 0.00009 | 14.21222 | Site_B | 12.90623 |
| 2 | 26 | 793.37631 | 0.31469 | 0.09539 | 0.00097 | 0.7181 | 0.19473 | 0.00023 | -0.02809 | 0.00738 | 0.00014 | 12.78199 | Site_B | 41.53379 |
| 5 | 26 | 771.74453 | 0.30826 | 0.09935 | 0.00192 | 1.50314 | 0.38825 | 0.00011 | -0.05683 | 0.0144 | 0.00008 | 13.22494 | Site_B | 19.90202 |
| 8 | 26 | 766.40058 | 0.29837 | 0.12221 | 0.01463 | 1.79784 | 0.68095 | 0.00829 | -0.06836 | 0.02545 | 0.00722 | 13.14974 | Site_B | 14.55806 |
| 11 | 26 | 756.39646 | 0.30137 | 0.10465 | 0.00398 | 2.28444 | 0.64861 | 0.00043 | -0.0866 | 0.02402 | 0.00031 | 13.18971 | Site_B | 4.55394 |
| 14 | 26 | 752.59519 | 0.29954 | 0.09878 | 0.00243 | 2.51336 | 0.58656 | 0.00002 | -0.09516 | 0.02154 | 0.00001 | 13.20555 | Site_B | 0.75267 |
| 17 | 26 | 752.48646 | 0.30092 | 0.09783 | 0.0021 | 2.61823 | 0.57445 | 0.00001 | -0.09874 | 0.0209 | 0 | 13.25772 | Site_B | 0.64394 |
| 20 | 26 | 751.84252 | 0.29872 | 0.09694 | 0.00206 | 2.85794 | 0.619 | 0 | -0.10646 | 0.02226 | 0 | 13.42245 | Site_B | 0 |
| 23 | 26 | 752.86719 | 0.29048 | 0.0971 | 0.00278 | 3.03013 | 0.65627 | 0 | -0.11153 | 0.02339 | 0 | 13.58468 | Site_B | 1.02467 |
| 26 | 26 | 755.8538 | 0.27848 | 0.09951 | 0.00514 | 3.09388 | 0.71927 | 0.00002 | -0.11235 | 0.0254 | 0.00001 | 13.76907 | Site_B | 4.01129 |
| 29 | 26 | 760.30966 | 0.27302 | 0.1037 | 0.00847 | 3.08653 | 0.78039 | 0.00008 | -0.11053 | 0.02729 | 0.00005 | 13.96219 | Site_B | 8.46715 |
| 32 | 26 | 762.04891 | 0.27426 | 0.10402 | 0.00837 | 3.17725 | 0.7859 | 0.00005 | -0.11267 | 0.02724 | 0.00004 | 14.09969 | Site_B | 10.20639 |
| 35 | 26 | 764.88911 | 0.27569 | 0.10372 | 0.00786 | 3.19872 | 0.78045 | 0.00004 | -0.11251 | 0.02682 | 0.00003 | 14.21504 | Site_B | 13.04659 |
| 2 | 29 | 796.60047 | 0.28726 | 0.10166 | 0.00472 | 0.70826 | 0.19812 | 0.00035 | -0.02769 | 0.00751 | 0.00023 | 12.78753 | Site_B | 44.75796 |
| 5 | 29 | 775.2205 | 0.27949 | 0.16131 | 0.08316 | 1.47824 | 0.79293 | 0.06228 | -0.0559 | 0.0296 | 0.05902 | 13.22336 | Site_B | 23.37798 |
| 8 | 29 | 769.50102 | 0.27867 | 0.08111 | 0.00059 | 1.77206 | 0.35512 | 0 | -0.0674 | 0.01302 | 0 | 13.14551 | Site_B | 17.6585 |
| 11 | 29 | 759.30379 | 0.28397 | 0.10045 | 0.0047 | 2.25278 | 0.44342 | 0 | -0.08543 | 0.01634 | 0 | 13.1845 | Site_B | 7.46127 |
| 14 | 29 | 755.20559 | 0.28438 | 0.13198 | 0.03119 | 2.48613 | 0.93981 | 0.00816 | -0.09421 | 0.03461 | 0.00649 | 13.19395 | Site_B | 3.36307 |
| 17 | 29 | 755.01947 | 0.28554 | 0.11287 | 0.01141 | 2.58842 | 0.71502 | 0.00029 | -0.09772 | 0.02605 | 0.00018 | 13.24467 | Site_B | 3.17695 |
| 20 | 29 | 754.22291 | 0.286 | 0.10799 | 0.00809 | 2.8343 | 0.71837 | 0.00008 | -0.10564 | 0.02583 | 0.00004 | 13.41513 | Site_B | 2.38039 |
| 23 | 29 | 754.83708 | 0.28402 | 0.10659 | 0.00771 | 3.03855 | 0.73779 | 0.00004 | -0.11185 | 0.02627 | 0.00002 | 13.58332 | Site_B | 2.99456 |
| 26 | 29 | 757.7127 | 0.27175 | 0.11259 | 0.01579 | 3.12163 | 0.87388 | 0.00035 | -0.11337 | 0.03087 | 0.00024 | 13.76724 | Site_B | 5.87019 |
| 29 | 29 | 762.33823 | 0.2603 | 0.1365 | 0.05654 | 3.11369 | 1.23331 | 0.01158 | -0.11153 | 0.04324 | 0.0099 | 13.95847 | Site_B | 10.49572 |
| 32 | 29 | 764.08342 | 0.25973 | 0.13196 | 0.04903 | 3.20435 | 1.17411 | 0.00635 | -0.11366 | 0.04082 | 0.00536 | 14.09656 | Site_B | 12.2409 |
| 35 | 29 | 766.97112 | 0.25946 | 0.12587 | 0.03927 | 3.22681 | 1.08871 | 0.00304 | -0.11351 | 0.03753 | 0.00249 | 14.21366 | Site_B | 15.1286 |
| 2 | 32 | 798.1306 | 0.27666 | 0.10735 | 0.00996 | 0.70985 | 0.20193 | 0.00044 | -0.02776 | 0.00767 | 0.00029 | 12.78693 | Site_B | 46.28808 |
| 5 | 32 | 777.18698 | 0.26149 | 0.08727 | 0.00273 | 1.46393 | 0.30401 | 0 | -0.0554 | 0.01117 | 0 | 13.21275 | Site_B | 25.34446 |
| 8 | 32 | 771.56283 | 0.26756 | 0.08821 | 0.00242 | 1.7488 | 0.35142 | 0 | -0.06657 | 0.01291 | 0 | 13.13594 | Site_B | 19.72031 |
| 11 | 32 | 761.23129 | 0.27406 | 0.08922 | 0.00213 | 2.22616 | 0.4266 | 0 | -0.08449 | 0.01557 | 0 | 13.17395 | Site_B | 9.38877 |
| 14 | 32 | 756.79787 | 0.27732 | 0.10887 | 0.01086 | 2.4705 | 0.47615 | 0 | -0.09368 | 0.01748 | 0 | 13.18598 | Site_B | 4.95535 |
| 17 | 32 | 756.41498 | 0.28157 | 0.1405 | 0.04507 | 2.57871 | 0.99275 | 0.00939 | -0.09743 | 0.0362 | 0.00712 | 13.2341 | Site_B | 4.57247 |
| 20 | 32 | 755.58367 | 0.28239 | 0.12195 | 0.02058 | 2.82273 | 0.84603 | 0.00085 | -0.10528 | 0.03042 | 0.00054 | 13.40544 | Site_B | 3.74115 |
| 23 | 32 | 756.01687 | 0.28384 | 0.11707 | 0.01533 | 3.04217 | 0.82171 | 0.00021 | -0.11202 | 0.02925 | 0.00013 | 13.57871 | Site_B | 4.17435 |
| 26 | 32 | 758.58101 | 0.27646 | 0.12614 | 0.0284 | 3.14987 | 1.02023 | 0.00202 | -0.11442 | 0.03603 | 0.00149 | 13.76417 | Site_B | 6.7385 |
| 29 | 32 | 763.2111 | 0.26268 | 0.27862 | 0.34579 | 3.14582 | 3.01223 | 0.29632 | -0.11273 | 0.10571 | 0.28626 | 13.95328 | Site_B | 11.36858 |
| 32 | 32 | 765.07214 | 0.25813 | 0.26271 | 0.32582 | 3.23383 | 2.84373 | 0.25546 | -0.11475 | 0.09905 | 0.24668 | 14.0913 | Site_B | 13.22962 |
| 35 | 32 | 767.9503 | 0.25701 | 0.17557 | 0.14324 | 3.25654 | 1.75587 | 0.06364 | -0.11459 | 0.06066 | 0.05888 | 14.21011 | Site_B | 16.10778 |
| 2 | 35 | 799.41947 | 0.26341 | 0.11089 | 0.01753 | 0.71518 | 0.20709 | 0.00055 | -0.02794 | 0.00786 | 0.00038 | 12.79774 | Site_B | 47.57695 |
| 5 | 35 | 778.42096 | 0.25321 | 0.09224 | 0.00605 | 1.46518 | 0.30357 | 0 | -0.05548 | 0.01117 | 0 | 13.20395 | Site_B | 26.57844 |
| 8 | 35 | 772.96678 | 0.25748 | 0.09375 | 0.00603 | 1.74171 | 0.34897 | 0 | -0.06635 | 0.01284 | 0 | 13.12474 | Site_B | 21.12426 |
| 11 | 35 | 762.76583 | 0.26356 | 0.09558 | 0.00583 | 2.21078 | 0.42248 | 0 | -0.08398 | 0.01545 | 0 | 13.16258 | Site_B | 10.92331 |
| 14 | 35 | 758.2436 | 0.2681 | 0.09606 | 0.00526 | 2.45412 | 0.45609 | 0 | -0.09313 | 0.01659 | 0 | 13.17559 | Site_B | 6.40108 |
| 17 | 35 | 757.71566 | 0.27202 | 0.1159 | 0.01893 | 2.5643 | 0.49491 | 0 | -0.09695 | 0.01805 | 0 | 13.22483 | Site_B | 5.87314 |
| 20 | 35 | 756.80521 | 0.27477 | 0.16673 | 0.09935 | 2.80904 | 1.41174 | 0.04662 | -0.10486 | 0.05079 | 0.03896 | 13.39468 | Site_B | 4.96269 |
| 23 | 35 | 757.24057 | 0.27587 | 0.13726 | 0.04445 | 3.02765 | 1.09436 | 0.00566 | -0.11156 | 0.03896 | 0.00419 | 13.5694 | Site_B | 5.39805 |
| 26 | 35 | 759.63684 | 0.27157 | 0.19342 | 0.16032 | 3.15111 | 1.99595 | 0.11439 | -0.11452 | 0.0705 | 0.1043 | 13.75809 | Site_B | 7.79432 |
| 29 | 35 | 764.05935 | 0.26057 | 0.11299 | 0.0211 | 3.16278 | 0.59974 | 0 | -0.11339 | 0.02092 | 0 | 13.94599 | Site_B | 12.21683 |
| 32 | 35 | 765.92986 | 0.25439 | 0.11255 | 0.02381 | 3.25339 | 0.61268 | 0 | -0.11552 | 0.02119 | 0 | 14.08204 | Site_B | 14.08735 |
| 35 | 35 | 768.90084 | 0.24993 | 0.112 | 0.02565 | 3.27467 | 0.61979 | 0 | -0.11529 | 0.02126 | 0 | 14.20148 | Site_B | 17.05832 |

